# An electrostatic cluster guides Aβ40 fibril formation in cerebral amyloid angiopathy

**DOI:** 10.1101/2022.12.22.521588

**Authors:** Elliot J. Crooks, Ziao Fu, Brandon A. Irizarry, Xiaoyue Zhu, William E. Van Nostrand, Saikat Chowdhury, Steven O. Smith

## Abstract

Cerebral amyloid angiopathy (CAA) is associated with the accumulation of fibrillar Aβ peptides upon and within the cerebral vasculature, which leads to loss of vascular integrity and contributes to disease progression in Alzheimer’s disease (AD). We investigate the structure of human-derived Aβ40 fibrils obtained from patients diagnosed with sporadic or familial Dutch-type (E22Q) CAA. Using cryo-EM, two primary structures are identified containing elements that have not been observed in in vitro Aβ40 fibril structures. One population has an ordered N-terminal fold comprised of two β-strands stabilized by electrostatic interactions involving D1, E22, D23 and K28. This charged cluster is disrupted in the second population, which exhibits a disordered N-terminus and is favored in fibrils derived from the familial Dutch-type CAA patient. These results illustrate differences between human-derived CAA and AD fibrils, and how familial CAA mutations guide fibril formation.

## INTRODUCTION

Cerebral vascular pathologies including cerebral amyloid angiopathy (CAA) are a major contributor to the progression of neurodegenerative diseases, occurring in 80% of Alzheimer’s Disease (AD) patients ^1,2^. Despite its prevalence, CAA remains largely untreatable and is often overlooked ^3^. The hallmark clinical presentations of CAA include vascular cognitive impairment and dementia, recurring intracerebral hemorrhage and stroke ^4^. Vascular deposition of the amyloid-β (Aβ) peptide upon and within the cerebral vasculature is the pathological trigger of CAA. Although often considered concomitantly as both share Aβ deposition as a hallmark, AD and CAA are distinct diseases that can occur independently ^5^.

Aβ peptides originate from non-specific proteolytic cleavage of C99, the β-C-terminal fragment of the amyloid precursor protein (APP) ^6^. The cleavage event results in peptides of multiple lengths, with the major product being Aβ40 ^7,8^, the principal isoform in vascular deposits ^9^. The earliest familial form of CAA identified was linked to a Dutch family ^10^ and originates from a glutamate to glutamine mutation at position 22 of the Aβ peptide ^11^. The resulting pathology, familial CAA Dutch (fCAA-Dutch), is an autosomal dominant form of vascular amyloidosis that is accompanied by accelerated cognitive impairment caused by extensive vascular amyloid deposition ^12^. fCAA-Dutch patients also spontaneously develop intracerebral hemorrhaging with the first stroke occurring on average at 25 years old ^10^. Pathologically, the disorder is characterized by CAA type-2 Aβ deposition in cortical and leptomeningeal small arteries and arterioles but in the absence of parenchymal cored amyloid plaques or neurofibrillary tangles that are key features of AD ^10,12^.

The Aβ peptide is highly polymorphic, adopting a wide array of conformations in solution ^13^. However, it is not clear whether the structural variability observed *in vitro* reflects the presence of polymorphism *in vivo* ^14^. The extent of structural variability of Aβ within a single patient remains debated. Some studies suggest one or two predominant structures exist per individual ^15–17^, while others find variability within a patient ^18^, or within amyloid plaques ^19^. Morphological differences between AD-specific parenchymal plaques and CAA-specific vascular deposits have been identified ^20–22^, suggesting a structural origin to the difference between the two pathologies. However, a robust link between structure and disease in the case of CAA has not been established, and the possibility of a structural origin to the variations in presentation, both within CAA and compared to AD, remains to be elucidated.

Clinical trials targeting Aβ have consistently underperformed. The poor conformational specificity resulting from the focus on *in vitro* fibrils or oligomers may have led to sub-optimal outcomes. Recent studies have demonstrated clear structural differences between amyloidogenic proteins incubated in solution and those extracted from human patients ^23^, including Aβ ^24–27^, and consequently motivate using brain-derived fibrils for structural studies. In addition, the lack of validated biomarkers for recognizing CAA and for differentiating CAA from AD ^28^ has been a major obstacle for clinicians.

Studies using electron cryo-microscopy (cryo-EM) ^24,27,29^ and solid-state nuclear magnetic resonance (NMR) spectroscopy ^15,26^ have investigated *ex vivo* Aβ fibril structures. Kollmer *et al*. (2019) ^24^ targeted vascular amyloid from cerebral meninges. They observed structural variability but were able to obtain a 4.4 Å structure of the predominant fibril population. The resulting structure revealed an ordered N-terminus and intermolecular hydrophobic packing, differing from structures of *in vitro* Aβ40 fibrils. The structure had some elements in common with fibrils seeded from parenchymal brain tissue using Aβ40 by Ghosh and colleagues ^29^, although the latter exhibited a disordered N-terminal segment. It is not clear whether these structures are specific to the individuals studied or characteristic of their respective disease states.

In this work, we compare the structures of Aβ40 fibrils seeded from vascular amyloid deposits isolated from sporadic CAA and fCAA-Dutch patients. We have recently shown that laser capture microdissection can be used to selectively isolate vascular amyloid deposits from brain slices ^30^. We use cryo-EM to determine fibril structures and identify multiple structural elements unique to brain vascular-derived fibrils, and use solid-state NMR spectroscopy to confirm the presence of these structural features. We find two structures primarily populate the patient-derived samples. Both structures exhibit a similar fold of the fibril core, characterized by an electrostatic cluster composed of residues E22-D23-K28. Importantly, the highly ordered N-terminal segment composed of two β-strands in one population is unique to *ex vivo* fibrils. A structured N-terminus is generally not observed in fibrils formed *in vitro*. However, the N-terminus of C99 is folded into a two-stranded β-hairpin prior to cleavage by γ-secretase. We show here that K28 stabilizes the structured N-terminus in both the γ-secretase substrate and brain-derived fibrils, highlighting the role of the electrostatic cluster in APP proteolytic processing and in fibril formation in the human brain. The comparison of the fibril structures from sporadic CAA and fCAA-Dutch patients highlights the importance of the E22-D23-K28 electrostatic cluster in fibril formation and in disease progression in CAA and AD.

## RESULTS

### Extraction and isolation of Aβ from patients

Vascular deposits containing Aβ fibrils were extracted from the cerebral vasculature of two patients, diagnosed with sporadic CAA and fCAA-Dutch. In contrast with other studies, we used laser capture microdissection to specifically target vascular amyloid ^30^. To further purify the samples and increase the length and number of Aβ fibrils, we employed templated fibril growth using wild-type Aβ40, which has been shown to propagate fibril conformation ^30,31^. Templated growth was verified using FTIR spectroscopy (Fig. S1) as described previously ^30^.

Morphology analyses were first performed on the cryo-EM datasets to characterize fibril homogeneity (Fig. 1 and S2). The Aβ peptide can exhibit extensive conformational variability that can hinder helical reconstruction in cryo-EM, and prior knowledge of the range of helical characteristics facilitates the separation of distinct fibril polymorphs. Morphology analyses revealed two primary fibril populations. The first population of fibrils (population A) exhibited narrow widths of ≈6.4 ± 0.9 nm and short crossover distances of ≈37 ± 3 nm. The second population of fibrils (population B) exhibited widths of ≈10 nm and a highly variable crossover distance averaging around 175 nm.

**Figure 1.**
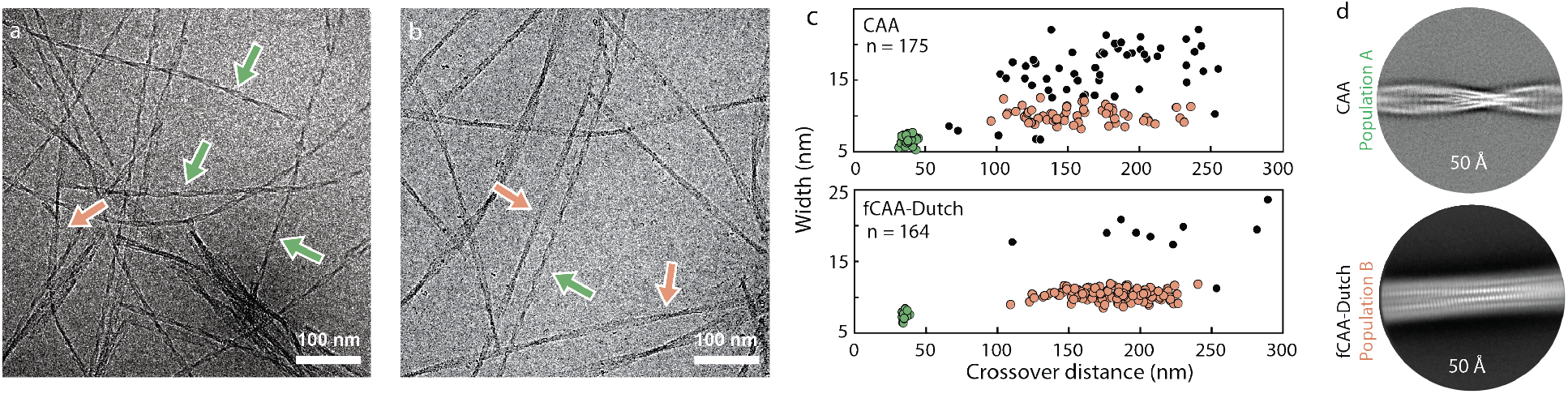
Two populations of Aβ fibrils in sporadic and familial Dutch vascular amyloid. Representative cryo-EM micrographs from the (a) CAA and (b) fCAA-Dutch patients. Arrows highlight different fibril populations. (c) Morphology plots for each patient. Each dot represents the average crossover-distance and width of a single fibril with colors corresponding to the arrows and different populations in panel (a). Two primary fibril clusters are observed: a highly twisted homogeneous population A (green), abundant in the CAA case, and a more heterogeneous and less-twisted population B (red), predominant in the fCAA-Dutch case. We note that the second population primarily exhibits variability in crossover distance but not in width, suggesting that the number of peptides in each cross-β unit is maintained along the fibril but the conformation may vary. The highly twisted fibril populations in both cases resulted in cryo-EM volumes with similar backbone traces. (d) 2D class averages from helical reconstruction reflect the difference in fibril morphology found in samples derived from sporadic and familial CAA patients. (top) The highly twisted fibrils abundant in the sample seeded from the CAA patient (in green in panel c) – named population A – have an average crossover distance of ≈37 nm. (bottom) The major fibril morphology in the sample seeded from the fCAA patient (red in panel c) exhibit a greater and more variable crossover distance.

The highly twisted fibril morphology was previously only observed in Aβ fibrils purified from the leptomeningeal vasculature of a CAA patient ^24^, whereas fibrils developed *in vitro* do not exhibit these characteristics, suggesting that this population is clinically relevant. The widths and longer crossover distances of population B, however, are not unique, being similar to most *in vitro* Aβ fibrils as well as fibrils seeded off parenchymal plaques of AD patients ^29^. On the basis of the morphology analyses, the highly twisted population A fibrils were found to be abundant in the CAA patient, while population B was the predominant fibril morphology in the fCAA-Dutch patient sample (Fig. S2). Both populations were targeted for helical reconstruction.

### Structure determination

We used cryo-EM to determine the structure of vascular amyloid fibrils seeded from both patients. From the two datasets, we obtained three cryo-EM maps (Fig. S3–S5). The population A structure was solved from data of the sporadic CAA patient to a resolution of 2.9 Å (Fig. 2a, S3). The population B structure was determined from data of the fCAA-Dutch patient to a resolution of 3.1 Å (Fig. 2b, S5). The high degree of morphological heterogeneity and low particle number of population B in the sporadic CAA patient sample precluded successful helical reconstruction. The low number of fibrils corresponding to population A in the fCAA-Dutch patient sample also hindered high-resolution reconstruction. However, using the population A structure as an initial model, we found that the same structure is likely present in the fibril pool from the fCAA-Dutch patient (Fig. S4).

**Figure 2.**
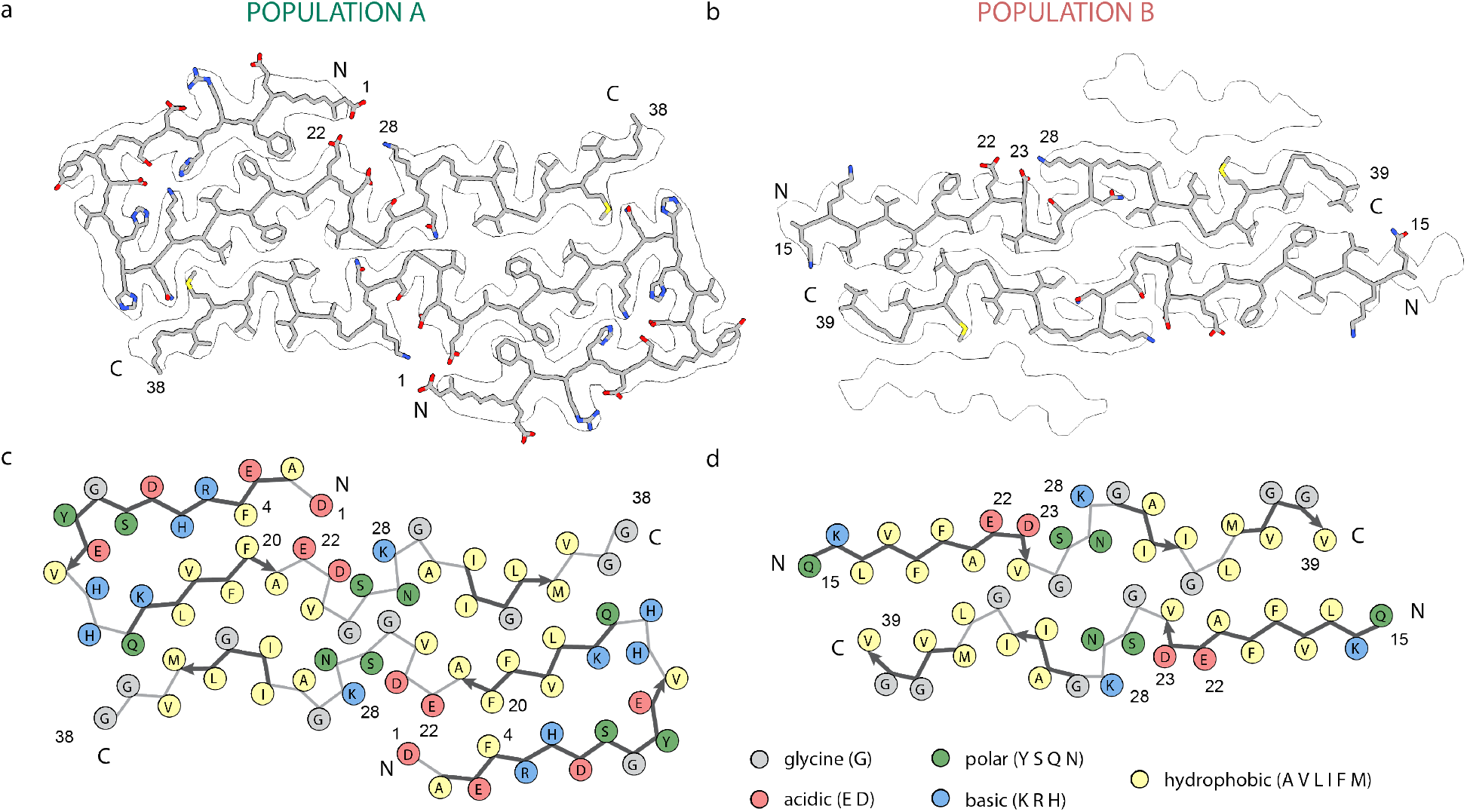
Cryo-EM Aβ40 structures from vascular amyloid of CAA patients. (a) Structure of population A fibrils. Nearly the entire Aβ40 monomer fits within the 2.9 Å cryo-EM density with well-defined positions for uncharged side chains. Population A is composed of two protofilaments with an ordered N-terminal section and inter-molecular hydrophobic interactions, with a β-strand from residues Q15 to A21. Reconstruction did not result in defined densities for the last two amino acids, V39 and V40. (b) Structure of population B fibrils within the outline of the cryo-EM density. Population B is composed of four layers. The 3.1 Å cryo-EM density identified a backbone fold with three β-strands composing each protofilament. The N-terminal region was not modeled due to a lack of resolvable density. (c) Schematic representation of population A with each residue colored according to its electrostatic property, as indicated in the legend. The two hydrophobic segments L17–A21 and A30–V36 interact in both structures but in an arrangement not found in many in vitro structures, suggesting a different mechanism of fibril formation. Multiple charged residue pairs are present in population A, both solvent-exposed and inside the fibril core. (d) Schematic representation of population B with each residue colored according to its electrostatic property as indicated in the legend. Similar to population A, hydrophobic segments L17–A21 and A30–V36 form intermolecular interactions. In this structure, charged residues do not form numerous interactions to stabilize the conformation.

The structure of population A is comprised of two cross β-units mediated by intermolecular hydrophobic interactions (Fig. 2a). In contrast with *in vitro* fibrils, the N-terminus has an ordered fold comprised of two β-strands mediated via several electrostatic interactions. The backbone fold of structure A is reminiscent of the conformation found by Kollmer *et al*. (2019) ^24^ (Fig. S6) Our model, however, differs in the form of a one-register shift, which we validate below.

The structure of population B fibrils is also formed from two cross β-units in the inner layers, but differs from population A in that the N-terminal segment remains disordered such that no defined density is visible prior to residue 14 (Fig. 2c). As in population A, the two hydrophobic segments interact in an inter-molecular fashion between protofilaments, distinguishing these structures from *in vitro* fibrils (Fig. 2d). We also found this population to exhibit a similar conformation to parenchymal-seeded Aβ40 fibrils determined by Ghosh *et al*. ^29^ from an AD patient (Fig. S6).

### Solid-state NMR spectroscopy as a probe of fibril structure and polymorphism

The structures of populations A and B were found to contain several elements typically not observed in *in vitro* fibrils, arguing that templated growth was able to capture aspects of fibril forms present in vascular deposits. Our structures also exhibit similarities and differences with Aβ fibril models previously developed from CAA and AD brain tissue. We used solid-state NMR spectroscopy to independently probe for specific structural features and assess polymorphism within the entire fibril population. We made use of several ^13^C labeled peptides, the first incorporating ^13^C labels at 1–^13^C F4, 2–^13^C L17, ring–^13^C F19, ring–^13^C F20, 2–^13^C G33, 5–^13^C M35, and 1–^13^C G38 (Fig. S7, S8).

We first investigated the unique two-stranded β-sheet fold of the N-terminus in population A by probing for a close F4-F20 contact (Fig. 3a), which is absent from *in vitro* fibrillar Aβ structures as well as published models of Aβ40 fibrils seeded from parenchymal amyloid ^15,29^ but present in *ex vivo* CAA fibrils ^24^. Evaluation of the resulting DARR NMR spectra reveals the presence of an F4–F20 cross-peak in samples from both the CAA and the fCAA-Dutch individuals where population A fibrils are present, thus supporting the existence of the N-terminal fold as determined via cryo-EM. The high resolution from this population also indicated a clear register shift starting at position 26 when compared to the structure determined by Kollmer *et al*. (2019) ^24^, resulting in a 180° flip of the C-terminal segment. We probed the L17–M35 contact specific to our model and observe a cross-peak (Fig. 3b) in both brain-derived samples, supporting the register shift and therefore the change in C-terminal orientation.

**Figure 3.**
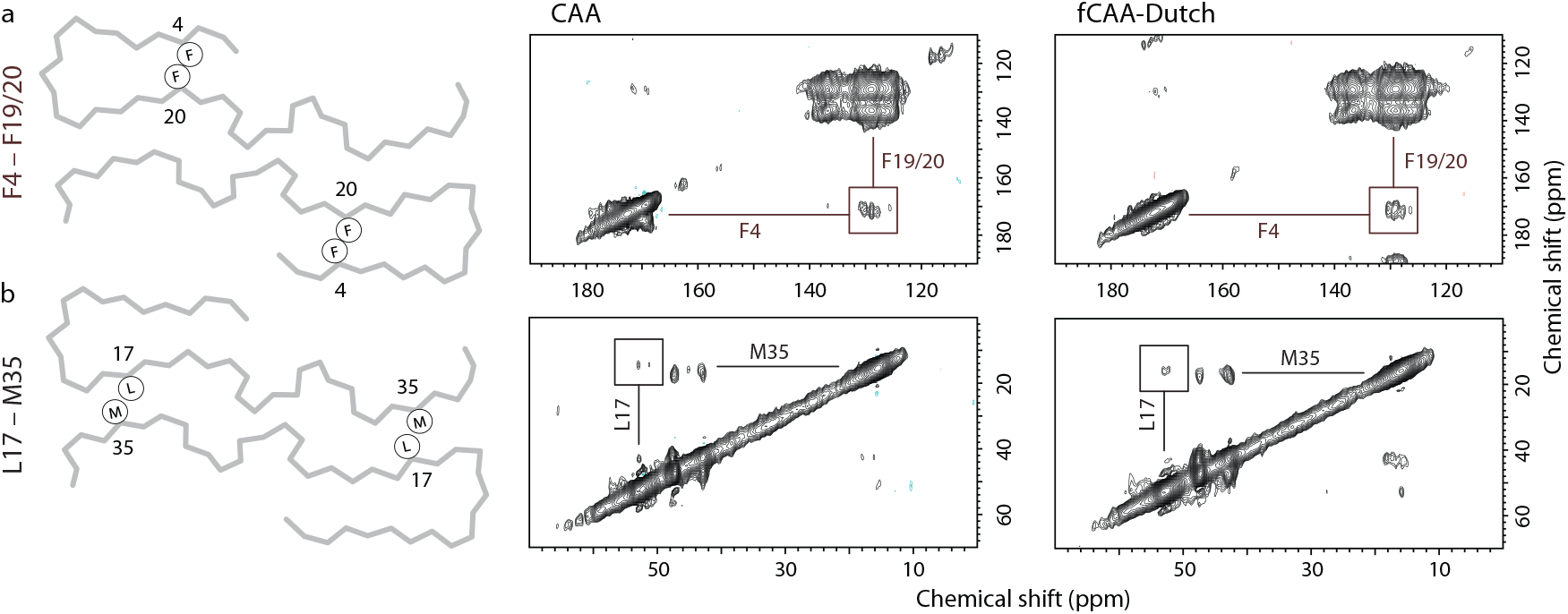
Probing brain-derived Aβ40 fibrils for population A with solid-state NMR. Two-dimensional ^13^C DARR NMR measurements of fibrils derived from sporadic CAA and fCAA-Dutch patients. The fibrils were produced via templated growth of brain amyloid using specifically ^13^C-labeled Aβ40 to probe inter-residue contacts modeled in the cryo-EM structures. (a) F4-F20 interaction. A scheme of the backbone trace of population A (left) highlighting the close F4-F20 intramolecular contact. Region of the 2D NMR spectrum (right) exhibiting cross peaks between 1-^13^C-F4 and ring-^13^C-F20 (or F19). The ^13^C labeling scheme chosen generally provides well-resolved resonances such that cross-peaks can be assigned to specific residue contacts. One exception is that the F19 and F20 ^13^C labels overlap but are predicted to interact with either G33 or F4, respectively. The presence of the F4-F19/20 cross peak (square) is consistent with the close F4-F20 distance in the cryo-EM structure in both the CAA and fCAA-Dutch samples. We also observe a cross peak between F19 and G33 (not shown), which is consistent with the population A structure. (b) L17-M35 interaction. A scheme of the backbone trace of population A (left) highlights the close L17-M35 intermolecular contact, while the 2D NMR spectrum (right) exhibits cross peaks between 2-^13^C-L17 and 5-^13^C-M35. The population A structure indicates a register shift with respect to the structure of Kollmer *et al*.^1^ leading to a flip of the C-terminal segment. This shift results in a close proximity between L17 and M35 consistent with the presence of observed cross-peaks in the NMR spectra of the CAA and fCAA-Dutch fibrils. The ^13^C chemical shifts taken from the cross peaks for the sCAA-sporadic fibrils are at 172.3 ± 0.3 ppm (1-^13^C-F4), 128.8 ± 0.2 ppm (1-^13^C-F19/F20), 50.7 ± 0.2 ppm (2-^13^C-L17) and 14.9 ± 0.3 ppm (5-^13^C-M35), and for fCAA-Dutch fibrils are at 172.3 ± 0.3 ppm (1-^13^C-F4), 128.6 ± 0.2 ppm (1-^13^C-F19/F20), 52.7 ± 0.2 ppm (1-^13^C-L17) and 15.3 ± 0.2 ppm (5-^13^C-M35).

The structure of population B is identical to fibrils seeded from parenchymal deposits of an AD patient ^29^ in that residues F19 and L34 are in close contact in the intermolecular interface (Fig. 2d). An F19-L34 interaction is also present in many structures of *in vitro* fibrils, also in an intermolecular fashion due to staggering of the individual monomers in the fibril, but with both hydrophobic residues in the interior of the fibril core. Using a second ^13^C-labeled Aβ peptide (ring–^13^C F19, 2-^13^C G33, U-^13^C L34, and 5-^13^C M35), NMR measurements confirm the presence of a close contact between the residues F19 and L34 (Fig. 4a). To investigate the intermolecular nature of this contact in structure B, we performed isotope dilution experiments and observe a decreased normalized cross-peak intensity consistent with the structure of population B (Fig. 4b).

**Figure 4.**
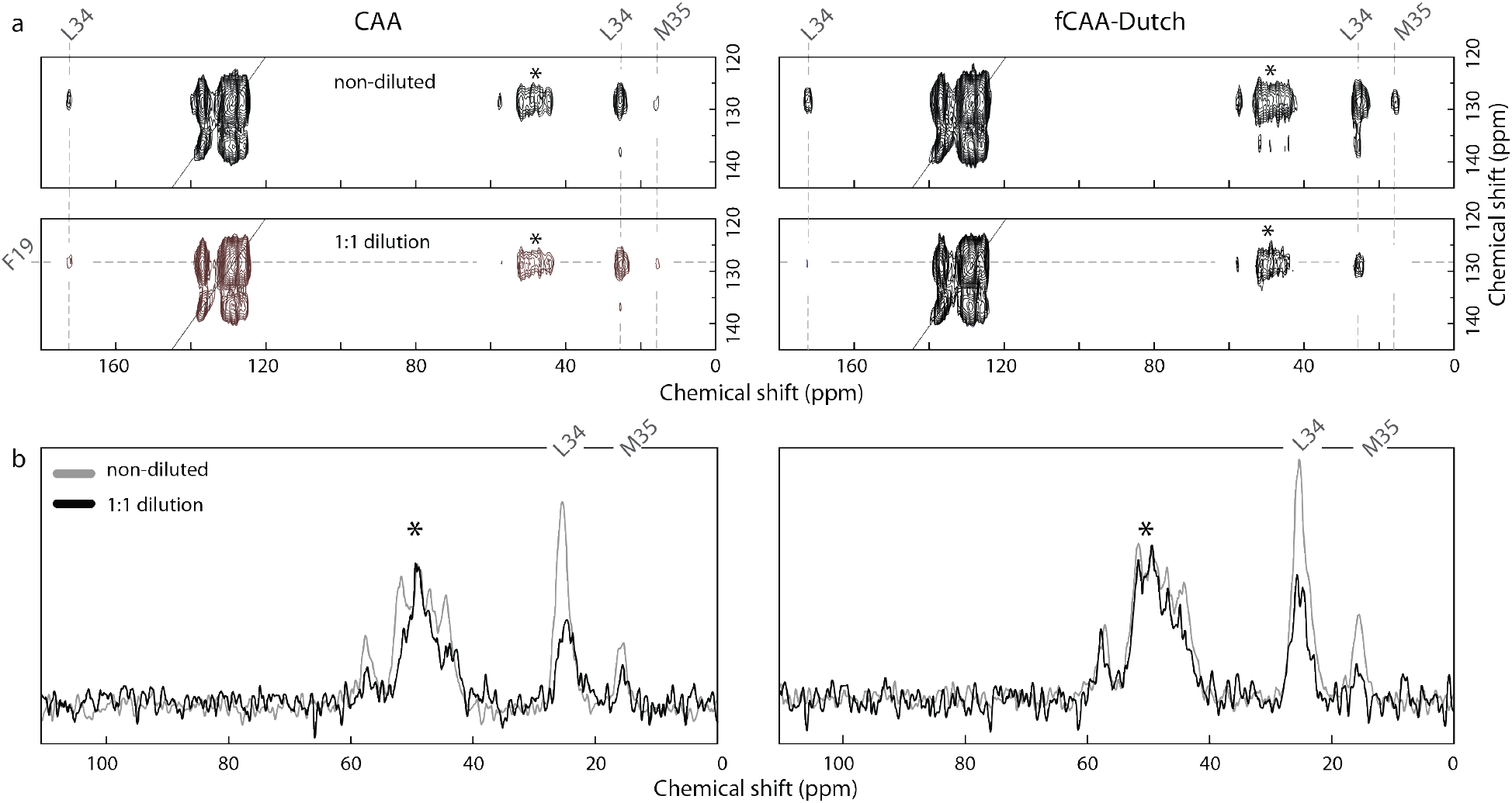
Probing brain-derived Aβ40 fibrils for population B with solid-state NMR. Two-dimensional ^13^C DARR NMR measurements of fibrils derived from sporadic CAA and fCAA patients to probe the contact between F19 and L34. The population B fibril structure is distinct from most *in vitro* fibrils by the linear shape of the inner Aβ peptides. This structure results in close proximity of F19 and L34 at the intermolecular interface. (a) Representative 2D ^13^C DARR NMR spectra are shown for the CAA and fCAA-Dutch patient samples after the third round of seeding (G3). (top) Region of the 2D NMR spectrum showing the diagonal resonance of ring-^13^C–Phe19 and crosspeaks to U–^13^C-Leu34 reflecting close Phe19-Leu34 packing in both CAA and fCAA-Dutch fibrils. (bottom) 1:1 isotope dilution indicate a marked intensity decrease of the ring-^13^C–Phe19 – U-^13^C–Leu34 crosspeaks in both patient samples. (b) Row through the diagonal ring-^13^C–Phe19 resonance showing cross peaks to Leu34. In order to assess whether the contacts are intra- or inter-molecular a parallel experiment was undertaken with ^13^C-labeled Aβ peptide diluted 50% with unlabeled Aβ peptide. Both samples exhibit a reduction in cross peak intensity consistent with inter-molecular interactions. Asterisks indicate MAS rotational side-bands. The ^13^C chemical shifts taken from the cross peaks for the sCAA-sporadic fibrils are at 25.8 ± 0.4 ppm (5-^13^C-L34), and 15.5 ± 0.3 ppm (5-^13^C-M35), and for the fCAA-Dutch fibrils are at 25.8 ± 0.3 ppm (5-^13^C-L34) and 15.5 ± 0.3 ppm (5-^13^C-M35).

The outer densities observed in the map of population B mirror findings from Ghosh *et al*. ^29^ who assigned these to β-hairpins. Although our map of population B suggests similar β-hairpins, the densities are too poorly defined to conclusively place these in this work. However, layering of these individual peptides atop the inner layers suggests a close intermolecular contact between residues F19 and M35, a contact that no other structure predicts. We observed the emergence of a cross-peak between these residues, consistent with β-hairpins populating these outer densities (Fig. S7).

### The E22-D23-K28 electrostatic cluster is a key structural element

The N-terminal structure in population A is unusual for Aβ fibrils. However, we have previously shown that the N-terminus adopts a two-stranded β-hairpin structure in C55, which contains the extracellular and transmembrane domains of C99 and releases Aβ upon proteolysis by γ-secretase ^32^. The observation of an ordered extracellular domain raises the possibility that the N-terminus in C99 influences Aβ fibril structure, and that the E22-D23-K28 electrostatic cluster is a key element for both C99 and Aβ fibrils.

Our previous studies on C55 and C99 indicate that substitutions in the KLVFF and YEV sequences strongly disrupted the N-terminal β-hairpin and resulted in increased Aβ production ^32^. Mutation of E22-D23, however, only slightly influenced the N-terminal β-hairpin structure and subsequent proteolysis ^32^, in contrast with K28 mutations that have been shown to influence γ-secretase processing. Specifically, K28A strongly decreased total Aβ production ^33^, while K28E increased total Aβ ^34^. To test whether the N-terminal β-hairpin structure in C55 extends to K28, we carried out FTIR measurements on C55 and the two K28 mutants, K28A and K28E. The amide I region in the FTIR spectrum is sensitive to secondary structure and contains two strong vibrational bands in wild-type C55: a 1655 cm^-1^ band, corresponding to the TM helix, and a 1627 cm^-1^ band, diagnostic of an N-terminal β-hairpin.

The K28A and K28E substitutions in C55 exhibit distinct behaviors, reflected primarily by changes in the intensity of the 1627 cm^-1^ band (Fig. 5a-c). This band becomes more pronounced in the K28A mutant indicating a stabilized N-terminal β-hairpin, but is lost in K28E–C55, consistent with unraveling the N-terminal structure. These changes argue that the interactions that stabilize the N-terminal β-hairpin in C55 extend to the E22-D23-K28 electrostatic cluster.

**Figure 5.**
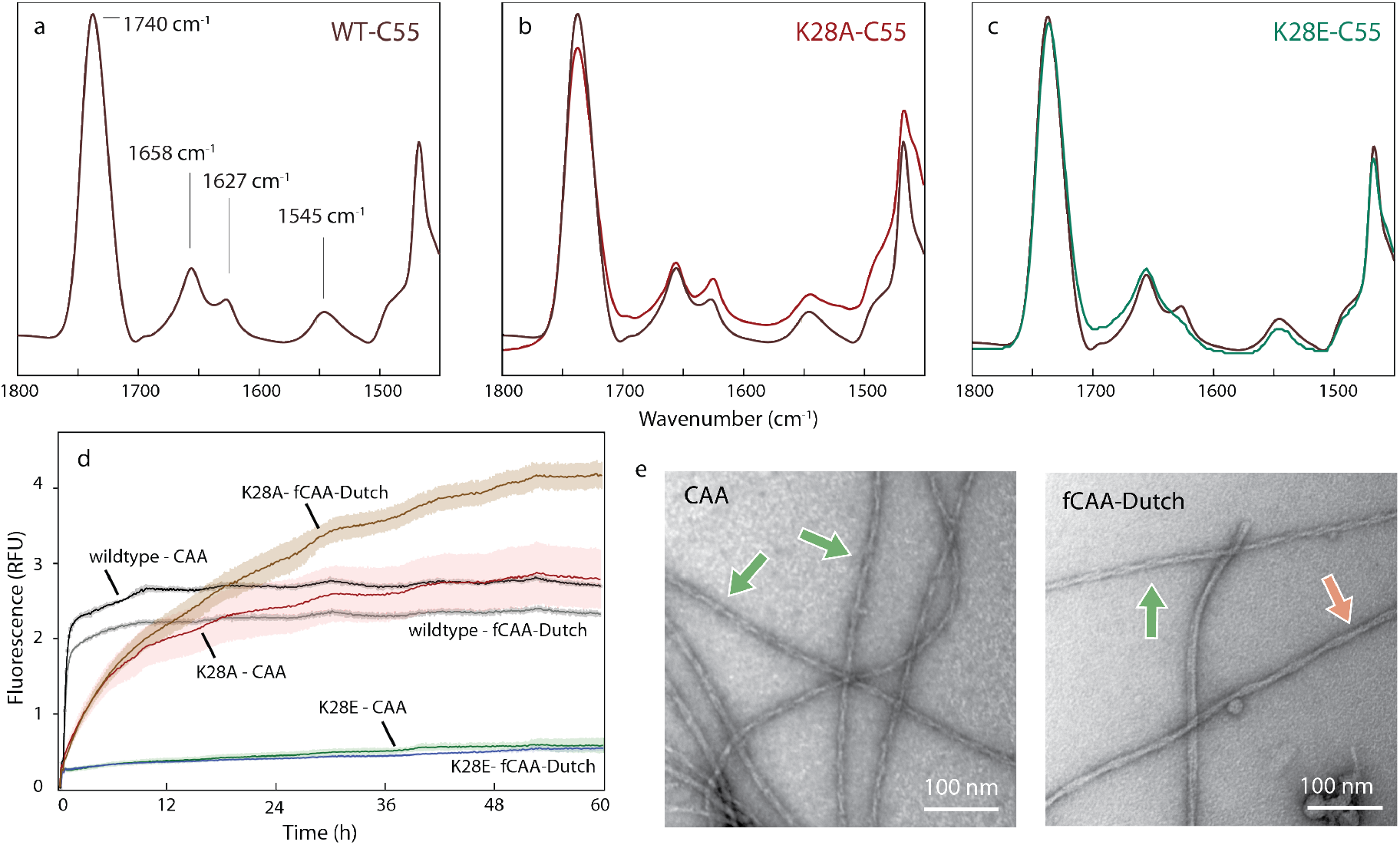
An electrostatic cluster mediates central interactions for both structures. Probing the E22-D23-K28 cluster in C55 and brain-derived Aβ40 fibrils. (a-c) FTIR spectra of (a) wild-type C55, (b) K28A C55 and (c) K28E C55 reconstituted into DMPC:DMPG bilayers. The acyl chain C=O vibration of the lipids is observed at ~1740 cm^-1^. The amide I region is between 1600 and 1700 cm^-1^ and the amide II region is between 1500-1560 cm^-1^. The band at ~1658 cm^-1^ corresponds to the TM α-helix, and the band at ~1627-1630 cm^-1^ corresponds to the N-terminal β-sheet. The K28A mutation increases the ~1630 cm^-1^ band relative to the 1658 cm^-1^ band, while the K28E mutation results in a decrease. (d) Templated growth upon G3 brain-derived fibrils was assessed by thioflavin T fluorescence using wild-type Aβ40, and the K28A and K28E mutants. For comparison, positive controls are shown (black and grey traces). The study was performed in triplicate and the results averaged, with noise representing standard deviation. The Aβ40 monomer containing the K28A mutation resulted in template growth for both brain-derived samples, but the initial fluorescence increase is slower than the positive control. The Aβ40 monomer containing the K28E mutation was not able to add to the brain-derived fibrils. (e) Representative negative stain TEM micrographs for the K28A Aβ40 mutant that successfully templated, as shown in (d), upon the CAA (top) and fCAA-Dutch (bottom) patient samples. We observe fibrils in all cases, with helical characteristics similar to population A (green arrow) or population B (red arrow).

We next investigated whether the K28 mutations influence the ability of the Aβ40 monomers to template off of the brain-seeded fibrils. In the population A fibril structure, the positively charged amine of K28 interacts with the negatively charged carboxylate of D23 in the neighboring monomer within the fibril (Fig. 6a). The cryo-EM map also reveals a second intermolecular electrostatic interaction between D1 and K28 (Fig. 6a). In this case, the K28 side chain on monomer *n* interacts with the D1 side chain of monomer *n+2* (Fig. 6b). This stagger may drive fibril elongation by acting as an anchor, locking in the added monomer. In contrast to population A, the N-terminus in the population B structure is not resolved (Fig. 2d). Here, K28 solely interacts with the negative carboxylate of D23 and no longer interacts with D1 (Fig. 6a). The D23-K28 interaction is intra-molecular and consequently the monomeric stagger observed for population A fibrils is absent, suggestive of a different mechanism of fibril elongation (Fig. 6c).

**Figure 6.**
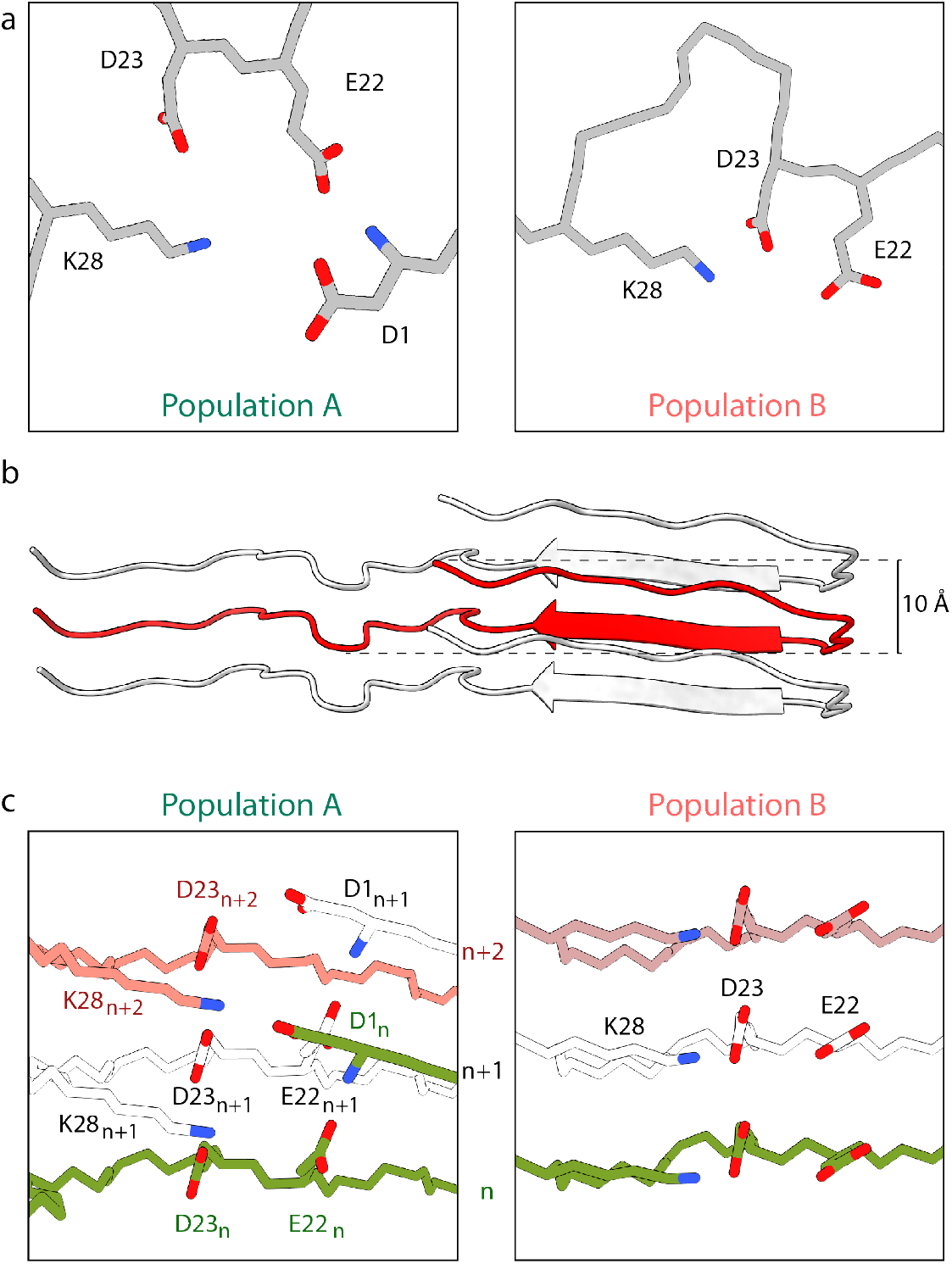
E22-D23-K28 cluster in populations A and B. (a) Top view of the E22-D23-K28 cluster in population A (left) and population B (right). In population A, charged residue pairs are brought into close proximity by the N-terminal fold, resulting in electrostatic interactions that stabilize and tether the N-terminus in position. Contrary to population A, the linear conformation of the inner layers in population B does not allow many electrostatic interactions. However, the D23–K28 salt bridge remains and is now able to be fully involved. (b) Ribbon view of population A highlights the out-of-plane displacement of a monomer in red up to 10 Å. (c) (left) Cartoon representation of population A with three individual monomers highlighting interactions between monomers (n), (n+1), and (n+2). Residues of interest are highlighted (colors represent different monomers). The N-terminus of monomer (n) (green) is displaced above the monomer plane and interacts with K28 two monomers above (n+2). Similarly, D23 interacts with K28 on the monomer (n+1). (right) In contrast, the same residues of the electrostatic cluster are interacting in an intermolecular fashion in population B, leading to the absence of stagger and inter-molecular stabilizing interactions.

We compared thioflavin T fluorescence measurements (Fig. 5d) to assess the ability of monomers of wild-type Aβ40 and the K28A-Aβ40 and K28E-Aβ40 mutants to undergo templated growth upon the brain-seeded fibrils. Wild-type Aβ40 efficiently polymerizes upon fibril seeds from the CAA and the fCAA-Dutch patients (Fig. 5d, S1). In contrast, the efficiency of K28A monomer addition to fibril seeds is reduced. TEM imaging of the K28A fibrils (Fig. 5e) indicate the presence of abundant, highly twisted fibrils, indicating that retaining the NH_3_^+^–E22 interaction may be sufficient to induce population A. In contrast to K28A, Aβ40-K28E monomers do not add to the *ex vivo* fibrils consistent with disruption of the E22-D23-K28 cluster and loss of N-terminal structure.

## DISCUSSION

We investigated the structure of Aβ40 fibrils seeded from vascular deposits of patients diagnosed with sporadic or familial Dutch-type (E22Q) CAA. Vascular amyloid deposits were extracted from brain tissue sections with laser capture microdissection and used for templated fibril growth. Using cryo-EM and NMR spectroscopy, we describe two structures, identified as populations A and B, which are markedly different from *in vitro* fibril structures. These share a common fibril core but differ in the fold of the N-terminal sequence. The structure of population A contains a well-folded N-terminus that is unique to brain-derived vascular fibrils and appears to originate from the two-stranded β-hairpin structure observed in the N-terminus of C55 and C99. Population B fibrils have a disordered N-terminus and are favored by the familial Dutch mutation. They are also observed in association with AD parenchymal deposits ^29^. We discuss below the potential importance of a structured N-terminus of Aβ40 fibrils and the central role of the E22-D23-K28 electrostatic cluster in guiding fibril formation in the human brain.

### Implications of a structured N-terminus in Aβ40 fibrils

The Aβ40 fibrils comprising population A appeared morphologically homogeneous and allowed cryo-EM helical reconstruction to a resolution of 2.9 Å (Fig. S3). The resulting structure exhibits parallel, in-register β-sheet and a well-folded N-terminal sequence. A similar conformation of the N-terminus was identified by Kollmer *et al*. (2019) ^24^ in fibrils isolated from leptomeningeal deposits of a sporadic CAA patient. The independent observation of similar N-terminal conformations in separate individuals, despite different extraction and purification methods, argues that the population A fibrils are a common feature of vascular amyloid.

The formation of a two-stranded β-sheet by the hydrophilic N-terminus stands in contrast with *in vitro* structures of wild-type Aβ40 and Aβ42. C99 also adopts a structured N-terminus prior to cleavage ^32^. In C99 (and C55) an N-terminal β-hairpin interacts with the surface of the lipid bilayer, enhancing electrostatic interactions, while the hydrophobic residues downstream of K28 form a membrane-spanning α-helix. Upon cleavage and release of Aβ from γ-secretase, the N-terminal two-stranded β-hairpin appears to convert to cross-β-sheet structure as Aβ monomers/oligomers polymerize into protofilaments en route to population A fibrils. The mature population A fibrils are then formed via inter-molecular packing of the hydrophobic L17-V18-F19-F20-A21 and C-terminal sequences of two protofilaments via inter-molecular interactions as observed in the sporadic CAA and fCAA-Dutch fibrils described here.

Several studies suggest that a structured N-terminus may be important in Aβ40 fibrils in human brain. The N-terminal Aβ(1–28) sequence can form fibrils in solution with a parallel, in-register geometry ^35^ that are stabilized by electrostatic interactions ^36^ indicating that deletion of the C-terminal hydrophobic stretch may allow the N-terminus to fold independently. Intrahippocampal injections of Aβ(1–28) into rats resulted in cognitive deficits ^37^, emphasizing the importance of the N-terminus in disease progression.

The observation of N-terminal β-sheet secondary structure provides an explanation to a long-standing question: Why are the α-cleavage products Aβ(17-40) and Aβ(17-42) non-amyloidogenic even though they contain both the Leu17–Ala21 and Ala30–Ala42 hydrophobic sequences? Our findings suggest that the N-terminus may be an important component of amyloid deposition and disease progression. Naturally occurring mutations offer evidence that N-terminal structure may influence phenotype. For instance, while the protective A2T mutation ^38^ decreases the hydrophobicity of the N-terminus, the A2V mutation ^39^ increases the amyloidogenesis and neurotoxicity of the mutated Aβ peptide. The increased hydrophobic character of the N-terminus increases the likelihood of β-sheet formation in the region, as evidenced computationally by reduced disorder and increased β-hairpin population in A2V–Aβ(1–28) ^40^. Furthermore, while Aβ42 is extensively modified post-translationally in the N-terminal region, Aβ40 exhibits significantly lower levels of modifications ^41^, indicative of the importance of the N-terminus in fibril and plaque formation in AD, and possibly in CAA.

Proteomics studies of patient brain extracts provide further evidence of an N-terminal fold *in vivo* ^41^. Aβ40–Aβ40 dimers, with D1–E22 as the primary cross-linkage, were found in patients with extensive levels of parenchymal Aβ40. These two residues interact in population A, and the formation of this crosslink may be explained by the co-localization of lysyl oxidase (LOX) in CAA-affected vessels ^42^. LOX is recruited to sites of brain injury ^43^, and converts primary amines (such as the N-terminal amine) into reactive aldehydes that can then form covalent bonds. A D1-E22 crosslink would stabilize the N-terminal fold observed in population A.

### The E22-D23-K28 electrostatic cluster guides fibril assembly

The E22-D23-K28 electrostatic cluster is an element common to both populations A and B. A bent backbone in this region was previously identified both in solution ^44,45^ and in some brain-derived fibrils ^29^, with a D23–K28 salt bridge present in many polymorphs ^44,45^. One of the key observations made in this work is that the N-terminal β-hairpin in the C55 substrate is influenced by the mutation of K28. The K28E mutation, which disrupts this structure, results in a dramatic increase in APP processing ^34^, similar to what was previously observed for mutation of the LVFF sequence ^32^. This observation provides additional evidence that the structured N-terminus in C55 and C99 is inhibitory and agrees with the abundance of shorter peptides resulting from α-secretase cleavage compared to the Aβ peptides resulting from β-secretase cleavage.

The E22-D23-K28 electrostatic cluster also provides an explanation as to why the structure of brain-derived fibrils differ from fibrils formed *in vitro*. Upon fibril formation, there appears to be a competition between the N- and C-termini to complex with the central region of the peptide. This idea has been proposed by Maji *et al*. ^46^, who found that a single D1Y substitution significantly altered Aβ fibril assembly. The observation of a similar E22-D23-K28 cluster in populations A and B suggests that loss of stabilizing D1-K28 and D1-E22 interactions and unraveling of the N-terminal fold in population A may be essential for exposing the electrostatic cluster and allowing binding of the β-hairpin structures in population B.

Mutations affecting the E22-D23-K28 interactions may influence a potential competition between the two fibril structures observed in vascular amyloid. Residues E22 and D23 within the cluster are the sites of several familial mutations that enhance vascular amyloid deposition, including the E22Q (Dutch), E22K (Italian) E22G (Arctic) and D23N (Iowa) mutations ^47^. All three familial mutations of E22 alter the electrostatic interactions of E22 with D1 and K28. Our results indicate that disruption of these interactions favors population B with the untethered N-terminus. However, the clinical presentation of each of these familial mutations is distinct from sporadic CAA. For example, the Italian E22K is associated with extensive cerebral vascular amyloid deposition, recurrent intracerebral hemorrhagic strokes, and the absence of neurofibrillary tangles reference ^48^. These clinical presentations are shared with fCAA-Dutch suggesting the Aβ40-Italian and Aβ40-Dutch peptides behave similarly in the cerebral vasculature, and may exhibit similar fibril structures.

Our studies on fibrils seeded from a fCAA-Dutch patient suggest that rather than generating new fibril forms, familial mutations may instead alter the ratio of different fibril populations. It should be noted, however, that we used the wild-type Aβ40 monomer to expand the fCAA-Dutch patient sample. The condition was chosen because the Aβ40-Dutch mutant peptide rapidly forms fibrils in solution, increasing the likelihood of forming *de novo* or self-nucleated fibrils. We have found that the wild-type Aβ40 and Aβ40-Dutch monomers are able to cross seed at comparable rates on fCAA-Dutch and sporadic CAA fibrils, respectively (unpublished results). As a result, using only the Aβ40 monomer for addition to fibril seeds eliminates self-nucleated fibrils, which do not form under our conditions for template growth ^30^. As discussed below, we might expect a further shift in the population ratios toward population B in the fCAA-Dutch case due to the higher propensity of the Aβ40-Dutch monomer to form β-hairpin structures that associate with the fibril surface in population B fibrils (see next section).

### Origins of fibril polymorphism *in vitro* and *in vivo*

The differences between brain-derived and *in vitro* fibril structures highlight the importance of the cellular environment in fibril formation ^14^. We discussed above the role of the existing N-terminal structure in C99 for guiding fibril formation in a cellular environment. Population A fibrils have well-defined widths and cross-over lengths, while population B fibrils exhibit much greater morphological variability. This structural polymorphism may originate from β-hairpins associated with the outer densities as described by Ghosh *et al*. ^29^, or from the association of metal ions or proteins to the N-terminus in cellular environments. There is a wide range of data supporting the rapid formation of β-hairpin intermediates on the pathway to Aβ fibrils ^49^, providing a potential explanation for their inclusion in population B. Monomeric Aβ spontaneously forms β-hairpins ^50^. However, we previously found that the Aβ40-Dutch peptide, and other fCAA mutants at positions E22 and D23, form β-hairpin structures much more rapidly than wild-type Aβ40 ^50^. The peptides may then layer upon pre-formed fibrils having the linear structure of population B. Moreover, the β-hairpins on the fibril surface may be responsible for the formation of secondary nucleation sites as the Dutch (E22Q) mutation, along with other fCAA mutations, increase secondary nucleation *in vitro* ^51^.

The differences in the N-terminal structures of populations A and B may also impact the ability to bind metal ions and other proteins, and result in fibril polymorphism. The N-terminus is generally regarded as a binding site for metal ions, where His6, His13, and His14 are known sites of coordination ^52^. These residues would only be available for coordinating metal ions in population B, which exhibits an unstructured N-terminus that coats the fibrils in a fuzzy, interactive sheath. Metal binding occurs in both vascular and parenchymal amyloid consistent with population B in both types of amyloids. In a similar fashion, an unstructured N-terminus might facilitate the interaction of other known binding partners such as clusterin and apolipoproteins ^53^.

### Implications for structure-disease correlations in amyloidopathies

Structure-disease correlations have been identified in other amyloidogenic diseases, such as ALS ^54^ and Parkinson’s disease ^55^, but have not been established in the case of AD or CAA. The possibility that the difference between AD and CAA originates at the structural level is supported by multiple recent findings ^20–22,56^. The results of this work add to this list; population A fibrils with an N-terminus comprised of two β-strands is unique to CAA. Our findings are supported by prior findings that N-terminal residues in vascular-derived fibrils are more ordered than in parenchyma-derived fibrils ^56^. As such, this study provides a high-resolution structure as a potential pharmacological target to distinguish CAA from AD. Recently, a novel β-hairpin fold of the N-terminus of Aβ has been used to generate antibodies that reduce plaque formation in mouse of AD ^57^, suggesting that the cross β-structure in the N-terminus of population A may provide a unique structural signature for antibody development. To date, antibodies developed toward AD appear to target an unstructured N-terminus. For example, the epitopes of aducanumab and gantenerumab target residues 3–7 and 2–9, respectively ^58,59^. These would be exposed in population B fibrils. An unstructured N-terminus is likely recognized by lecanemab, a therapeutic antibody that has recently yielded significant improvements in reducing cognitive decline in a phase three clinical trial, although with significant side effects involving brain edema and hemorrhaging ^60^. The observation that population B fibrils are identical to fibrils found in parenchymal amyloid suggests that lecanemab binding to vascular amyloid results in amyloid disaggregation and disruption of the blood vessel walls leading to hemorrhaging.

Finally, the differences in fibril populations between sporadic and fCAA-Dutch may be associated with the differences in clinical phenotype. The clinical presentation of fCAA-Dutch involves fewer ischemic strokes and more immune-active vascular amyloid deposits than sporadic CAA ^10,11,47^. Further structural studies of fibrils derived from vascular amyloid, in conjunction with proteomics studies to assess differences in metal and protein interactions, will be valuable in addressing the origins of the different phenotypes that have been described for CAA.

## Acknowledgements

This work was supported through grants from the National Institutes of Health to SOS and WVN (AG-027317 and AG-061775). Cryo-EM data were collected at the Stony Brook University cryo-EM center, which was established with National Institutes of Health Grant number S10 OD012272. We thank Drs. Sjors Scheres and Takanori Nakane for their useful help and suggestions with cryo-EM data processing, and Drs. Kent Thurber and Robert Tycko for their MATLAB scripts for optimizing particle alignment. SOS is grateful to Dr. Lucia Alvarez-Gutierrez and Matthias Koch for discussions and insights into the importance of the E22-D23-K28 cluster in the processing of C99.

## Ethics declaration (competing interests)

The authors declare no competing interests.

## Author contributions

E.J.C. and S.O.S. conceived the study and wrote the manuscript. E.J.C. performed most of the experiments. X.Z. and W.V.N. isolated vascular amyloid by laser capture microdissection. B.A.I. generated fibrils via seeded growth from brain amyloid and performed some spectroscopy experiments. Z.F. initiated the cryo-EM component of project and assisted with the atomic model building in the cryo-EM maps. S.C. assisted in helical reconstruction and cryo-EM data collection.

## METHODS

### Synthetic Aβ40

Wild-type, mutant, and isotope-edited amyloid-β peptides were synthesized using tBOC-chemistry ERI-Amyloid (Waterbury, CT) and purified by high-performance liquid chromatography using linear water–acetonitrile gradients containing 0.1% (v/v) trifluoroacetic acid. The mass of the purified peptide was measured using matrix-assisted laser desorption or electrospray ionization mass spectrometry and was consistent with the calculated mass for the peptide. On the basis of analytical reverse-phase high-performance liquid chromatography and mass spectrometry, the purity of the peptides was 95–99%.

### Isolation and templated growth of cerebral vascular amyloid deposits from human brain

Tissues used in these studies were from a 71-year-old male sporadic CAA patient (University of California Irvine Alzheimer’s Disease Research Center) and a 76-year-old familial CAA-Dutch patient (Leiden University Medical Center, Leiden, NL). The samples used here were prepared as described in Irizarry *et al*. ^30^. Briefly, for each case, individual amyloid-containing cerebral vessels were identified, excised and captured using a LMD6 laser capture microdissection microscope LMD6 (Leica Microsystems). The dissected vascular amyloid deposits were collected into sodium phosphate buffer, pH 7.2. Three rounds of templated growth were then performed using synthetic wild-type amyloid-β (1–40) at 15% seed content in sodium phosphate buffer.

### Fourier-transformed infrared spectroscopy

FTIR measurements were made with a Bruker Vertex 70v spectrometer with a room temperature detector and attenuated total reflectance (ATR) accessory. Samples were layered on a 2 mm germanium ATR plate (Pike Technologies) by drying 100 μL of peptide sample on the Ge surface with a stream of air. The spectral resolution was 4 cm^-1^.

The C55 peptides were co-solubilized in DMPC, DMPG, and octyl-β-glucoside in hexafluoroisopropanol. The molar ratio of peptide:lipid was 1:60 and the molar ratio of DMPC:DMPG was 10:3. The solution was incubated overnight at 37 °C, after which the solvent was removed under a stream of argon gas. The dried mixture was rehydrated in HEPES buffer (10 mM HEPES, 50 mM NaCl, pH 7.0) gently mixed at 37 °C for 6 h. The octyl-ß-glucoside (2% w/v) was removed by dialysis using Spectra-Por dialysis tubing with a 3500 MW cutoff.

### Nuclear magnetic resonance spectroscopy

Room temperature solid-state magic angle spinning (MAS) NMR experiments were performed at a ^13^C frequency of 125 MHz on a Bruker AVANCE spectrometer using 4 mm MAS probes. The MAS spinning rate was set to either 9 or 12 kHz to prevent rotational sidebands from covering target cross-peaks. Ramped amplitude cross polarization was used with a contact time of 2 ms. The ^13^C field strength was 54.4 kHz and ramped ^1^H field was centered at approximately 50 kHz. Two-pulse phase-modulated decoupling was used during the evolution and acquisition periods with a radiofrequency field strength of 82.7 kHz. Internuclear ^13^C–^13^C distance constraints were obtained from 2D dipolar assisted rotational resonance (DARR) NMR experiments using a mixing time of 600 ms. Each data set contained 64 t1 increments and 1024 complex t_2_ points with spectral widths of 27.7 kHz in both dimensions. 512 scans were averaged per t_1_ increment. All ^13^C solid-state MAS NMR spectra were externally referenced to the ^13^C resonance of neat TMS at 0 ppm at room temperature. Using TMS as the external reference, we calibrated the carbonyl resonance of solid glycine at 176.46 ppm. The chemical shift difference between ^13^C of DSS in D_2_O relative to neat TMS is 2.01 ppm.

Samples were prepared via templated growth as described in Irizarry *et al*. ^30^ using ^13^C-labeled peptides. The sample used as the template was the one used for cryo-EM acquisition, except for the NMR experiments using the ring–^13^C F19, 2-^13^C G33, U-^13^C L34, 5-^13^C M35 labeled peptide for the CAA patient (shown in Fig. 4), which used generation 5 fibrils as the template.

### Thioflavin T fluorescence spectroscopy

Fluorescence measurements were taken using a Spectra Max iD3 spectrometer (Molecular Devices). A final concentration of 37.5 μM thioflavin-T was used with an excitation wavelength of 440 nm and an emission wavelength of 490 nm in a 96-well clear Greiner microplate. Measurements were taken every 10 min for 80 h with 2 s of low orbital shaking in-between reads. The OD setting was set to 1. Negative blank and thioflavin T controls were included. All conditions were replicated three times, and the results averaged with standard deviation used to assess error. The overall experiments were repeated twice to test reproducibility; we show only one iteration.

The fluorescence kinetic experiments used samples prepared via templated growth as described in Irizarry *et al*. ^30^ but using a 10% seed content to enhance potential differences in efficiency.

### Negative stain transmission electron microscopy

Samples were diluted, deposited onto freshly glow-discharged carbon-coated copper 400 mesh grids (Electron Microscopy Sciences). Excess sample and buffer were removed by wicking with a Whatman 1 filter paper and immediately followed by negative staining with 2% (w/v) freshly filtered uranyl formate solution and air dried. The samples were imaged on a FEI Tecnai 12 BioTwin 80 kV transmission electron microscope with micrographs captured with an Advanced Microscopy Techniques camera at nominal magnification of 250,000x, corresponding to 2 Å/pixel.

### Cryo-electron microscopy

Following two-fold dilution in deionized water, 3.5 μL of sample were applied on a glow-discharged holey gold grid (UltrAuFoil Au R1.2/1.3, 300 mesh, Electron Microscopy Sciences), blotted for 3 seconds, at 4 °C and 100% relative humidity, followed by plunge freezing in liquid ethane (at −186°C) using an FEI Vitrobot Mark IV robotic plunge freezing device.

Data acquisition for cryo-EM was performed on a 200kV Talos Arctica (FEI, Thermo Fisher Scientific) transmission electron microscope equipped with a Falcon 3EC direct electron detector (FEI, Thermo Fisher Scientific). Imaging for both CAA and fCAA-Dutch samples were performed as movies (100 fractions) per micrograph in electron counting mode at a nominal magnification of 92,000x, corresponding to a detector pixel size of 1.12 Å/pixel. Micrographs for CAA samples were collected with under focus values varying between 0.8-1.4 μm and for the fCAA-Dutch samples between 0.2-1.4 μm. The total exposure dose for each micrograph for CAA and fCAA-Dutch samples were 49.7 e-/Å^2^ and 56.92 e-/Å^2^ respectively. The parameters for cryo-EM acquisition and helical reconstruction can be found in Table 1 (Extended Data) and the atomic model validation statistics can be found in Table 2 (Extended Data).

**Table 1.** Cryo-EM acquisition and helical reconstruction.

|  | CAA | fCAA-Dutch |  |
| --- | --- | --- | --- |
| Data collection |  |  |  |
| Microscope | Talos Arctica | Talos Arctica |  |
| Detector | Falcon III | Falcon III |  |
| Magnification | 92,000 | 92,000 |  |
| Voltage (kV) | 200 | 200 |  |
| Electron exposure (e<sup>-</sup>/Å<sup>2</sup>) | 49.70 | 56.92 |  |
| Electron flux (e<sup>-</sup>/pixel/sec) | 0.72 | 0.82 |  |
| Defocus range (μm) | 0.8–1.4 | 0.6–1.4 |  |
| Pixel size (Å) | 1.12 | 1.12 |  |
| Total exposure time (sec) | 85.83 | 86.11 |  |
| Fractions/micrographs | 100 | 100 |  |
| Exposure per fraction (e<sup>-</sup>/Å<sup>2</sup>/frame) | 0.4970 | 0.5692 |  |
| Micrographs collected | 2075 | 2140 |  |
| Processing |  |  |  |
|  | Population A | Population A | Population B |
| Symmetry imposed | 2<sub>1</sub>-screw | 2<sub>1</sub>-screw | 2<sub>1</sub>-screw |
| Rise (Å) | 2.44 | 2.41 | 2.44 |
| Twist (°) | 181.17 | 181.17 | 179.75 |
| Initial particle images (no.) | 68,351 | 4,107 | 114,106 |
| Final particle images (no.) | 59,075 | 4,081 | 73,287 |
| Map resolution (Å) | 2.87 | 7.77 | 3.09 |
| FSC threshold | 0.143 | 0.143 | 0.143 |
| *B*-factor (Å<sup>2</sup>) | -31.1 |  | -81.5 |
| EMDB ID | 29036 |  | 29037 |

**Table 2.** Atomic model validation statistics.

|  | CAA | fCAA-Dutch |
| --- | --- | --- |
|  | Population A | Population B<br>Inner layers only |
| Initial model used (PDB code) | 6SHS | 6W0O |
| **Model composition** |  |  |
| Non-hydrogen atoms | 2,910 | 1,098 |
| Protein residues | 380 | 150 |
| **Refinement** |  |  |
| Refinement package | Phenix<sup>3</sup> | Phenix<sup>3</sup> |
| CC (mask) | 0.8 | 0.67 |
| **R.m.s. deviations** |  |  |
| Bond lengths (Å) | 0.005 | 0.002 |
| Bond angles (°) | 0.427 | 0.432 |
| **Validation** |  |  |
| EMRinger<sup>4</sup> score | 6.45 | 4.94 |
| MolProbity<sup>5</sup> score | 1.26 | 0.94 |
| Clashscore | 4.96 | 1.83 |
| Poor rotamers (%) | 0 | 0 |
| CaBLAM outliers (%) | 0 | 0.79 |
| Cβ deviations (%) | 0 | 0 |
| Ramachandran |  |  |
| Favored (%) | 100 | 100 |
| Allowed (%) | 0 | 0 |
| Disallowed (%) | 0 | 0 |
| **Deposition** |  |  |
| PDB | 8FF2 | 8FF3 |
| EMD | 29036 | 29037 |

### Morphology assay

Motion-corrected micrographs (as detailed below) were converted to JPEG format using the mrc2tif program from the IMOD package ^61^. A Gaussian blur was applied, and the contrast was enhanced for better visualization using ImageJ software. For all fibrils clearly distinguishable, i.e., unobstructed by intra-fibrillar interactions, both the width and cross-over distances at multiple points along each fibril were measured. Using an in-home written MATLAB script, average width and cross-over distance per fibril was calculated and plotted. The script can be found on GitHub (*https://github.com/*) under *elliotcrooks/fibril. git*.

## Data availability

Density maps for the CAA (population A) and fCAA-Dutch (population B) samples were deposited in the EM Data Bank with reference IDs EMD-29036 and EMD-29037, respectively, and corresponding atomic models deposited in the Protein Data Bank (PDB) with reference IDs 8FF2 and 8FF3, respectively. The density map for population A in the fCAA-Dutch sample was deposited in the EM Data Bank with reference ID EMD-29038. NMR chemical shifts are reported in the figure legends. Raw 2D NMR data sets collected at the Keck NMR Center for Structure Biology are available from the corresponding author (SOS) upon request. The authors declare that all other data supporting the findings of this study are available within the article and its supplementary information files.

## Code availability

A MATLAB script was used in part of the data analysis. The script utilizes length measurements acquired manually in ImageJ. Each unique filament is separated by a measurement of small value by the user. The script then segments each filament and extracts averages of fibril characteristics. The script can be found on GitHub at *https://github.com/elliotcrooks/fibril*.

## EXTENDED CRYO-EM PROCEDURES

For all datasets, movie frames were motion-corrected and dose-weighed using MotionCor2 ^62^ with an applied B-factor of 500 and 8×8 patches. Aligned and dose-weighed micrographs were used to estimate contrast transfer function (CTF) using Gctf ^63^, with local refinement and validation. All subsequent image processing steps were performed in RELION 3.1 ^64^.

### Sporadic CAA/AD: patient population A

#### Helical reconstruction

Helical processing was carried out using RELION ^64^. Filaments were picked manually and extracted with a box size of 500 pixels and an inter-box distance of 47.5 Å. Multiple (4) rounds of reference-free 2D classification (K=100, T=1) were performed, discarding particles contributing to suboptimal 2D averages. Helical offset was left unrestricted and bimodal angular searches were allowed. We found that allowing the ‘Ignore CTF until the first peak’ function in the penultimate round of 2D classification allowed for better particle curation. A featureless cylinder was used as the initial model using the RELION_helix_toolbox program, with a 90% spherical mask applied. The diameter of the cylinder was determined from the morphology assay described above. Helical parameters (twist and rise) were estimated from the morphology assay and the FFT of 2D class averages exhibiting clear β-sheet separation. Two successive rounds of 3D classification with a single class requested, a regularization parameter T of 3, and applied C2 helical symmetry was first used for broad alignment. The resulting volume showed β-sheet separation and was low-passed filtered to 10 Å for subsequent 3D autorefinement, with optimization of helical rise and twist. The local averaging feature, as initially described by Ghosh *et al*. ^29^, was implemented to allow orientational and translational assignments to be consistent between segments of each fibril. Briefly, the feature uses the distance between particles along a given fibril and attempts to adjust orientational assignments such that they agree with the expected pitch. In addition, correction of the Fourier-shell correlation (FSC) curves (with phase-randomization) using masked half-maps at each iteration was found to be crucial in reaching convergence and was applied, using the solvent-flatted FSC function, for all 3D auto-refinements hereafter. Resolution at this stage reached 4.2 Å, with helical parameters of ±2.23° twist and 4.82 Å rise. We then performed Bayesian polishing followed by auto-refinement, lowering the reference low-pass filtering to 7 Å and allowing helical rise and twist optimization. CTF refinement procedures were performed first for beam-tilt estimation, then (anisotropic) magnification, followed by per-particle defocus fitting. Polished and CTF-corrected particles were used for 3D auto-refinement without helical parameter optimization, resulting in a 3.1 Å resolution volume. Upon post-processing with RELION, we found some residual misaligned particles or overfitting and used 3D classification (T=3, K=2) with helical parameter optimization to separate those particles, which did not contribute positively to the reconstruction. The most populated class was selected for an additional round of 3D auto-refinement, Bayesian polishing, CTF refinement, and 3D auto-refinement, resulting in a resolution estimate of 3.1 Å.

Upon inspection of the final volume and 2D class averages, and in agreement with previous studies on *ex vivo* amyloid-β ^24,29^, we identified possible 2_1_ screw helical symmetry. We performed 3D auto-refinement using the 3.1 Å volume, low-pass filtered to 4.5 Å to retain β-strand separation, as initial model, with an applied 2_1_ screw symmetry and allowing helical parameters to modulate. Half of the optimized helical rise was used for initial input, and (360 – r)/2 for initial twist, r being the optimized rise of the previous reconstruction. It should be noted that cryo-EM does not provide any information of helical filament handedness. As such, prior knowledge is required. In our case, right-handedness was assumed based on prior studies ^24^. Another subsequent refinement with symmetry search surpassed the 3.0 Å resolution threshold, with parameters of 2.41 Å (rise) and 181.31° (twist). The following corrections were then applied: CTF refinements (third order aberration than per-particle defocus fitting) followed by refinement, then CTF refinements as above followed by Bayesian polishing (omitting the final 20 frames to avoid radiation-induced motions) and auto-refinement. A final round of CTF corrections was performed, first estimating magnification anisotropy—which was negligible—, then third and fourth order aberrations, then per-particle defocus and per-micrograph astigmatism. We note in particular that avoiding per-micrograph B-factor estimation was crucial in preventing the introduction of errors. The final volume (59, 075 particles) reached a resolution of 2.9 Å as estimated from an FSC of 0.143, with 2_1_-screw symmetry, a rise of 2.44 Å and a twist of 181.17°. This final reconstruction was used for map sharpening and model building (Figure S3).

#### Map sharpening and model building

Post-refinement map sharpening was performed using deepEMhancer ^65^ using half-maps from the final refinement. User-provided noise statistics instead of automatic estimation of normalization parameters performed better sharpening of the map.

An Aβ atomic model 6SHS ^24^ was used as the initial model. Aβ monomers were individually rigid body fitted into the map using UCSF Chimera ^66^. We found a register shift starting at position 26 was required for the sidechains to fit properly into the density. This and other residues were manually fixed in Coot ^67^ with local real space refinement. The model was then submitted to multiple rounds of real-space refinements in Phenix ^68^. Due to missing map density, residues V39 and V40 could not be modeled. These were added post-hoc for the figures above but omitted from the final atomic model. After manually adjusting the model in Coot ^67^ and ISOLDE ^69^ to alleviate clashes, a final real-space refinement was performed with secondary structure restraints and without morphing or simulated annealing. MolProbity ^70^ and EMRinger ^71^ were used for model validation.

### fCAA-Dutch patient: population A

#### Helical reconstruction

Filaments were picked manually and extracted with a box size of 500 pixels and an inter-box distance of 48.3 Å in RELION, resulting in a ≈4,000 particles stack. Two rounds of reference-free 2D classification (K=50, T=2) were performed, with few discarded particles.

3D auto-refinement procedures were undertaken using the final volume from population A of the CAA patient sample as the initial model. The choice was made as an initial refinement procedure with a featureless cylinder suggested a similar structure as the one found in the sporadic CAA/AD patient sample. The low-pass filter cut-off for the initial model was purposefully kept at 4 Å to allow the β-strand separation to guide refinement calculations. Symmetry parameters refined in the prior reconstruction were used here as well. We find again that FSC correction using masked half-maps after each iteration helped ameliorate results. The final volume reached an estimated resolution around 7 Å with notably high B-factor, reflecting the difficulty of aligning a small particle stack. It should be noted here that further per-particle corrections (motion or aberrations) of a small particle stack will likely introduce biasing issues. However, the small particle number knowingly could not produce a high-resolution map. Nonetheless, we proceeded with Bayesian polishing, CTF refinement, and subsequent 3D auto-refinement in the same manner described above. The final volume harbored a resolution around 7 Å as estimated from a FSC of 0.143 (Figure S4), with overfitting and large B-factor. An increased correlation around 4.7 Å is observed and reflects the bias introduced by the initial model used as expected. Map sharpening or atomic model fitting procedures were not undertaken due to low resolution and inappropriate estimation of resolution.

### fCAA-Dutch patient: population B

#### Helical reconstruction

Filaments were picked manually and extracted with a box size of 250 pixels and an inter-box distance of 48 Å. Multiple (4) rounds of reference-free 2D classification (K=100, T=2) were performed, discarding particles contributing to suboptimal 2D averages. Helical offset was left unrestricted and bimodal angular searches were allowed, as polarity along the helical axis was assumed from prior studies. The ‘Ignore CTF until the first peak’ function was applied in the penultimate round of 2D classification. A featureless cylinder was used as initial model using the RELION_helix_toolbox program, with a 90% spherical mask applied. The diameter of the cylinder was determined from the morphology assay described above. Helical parameters (twist and rise) were estimated from the morphology assay and the FFT of 2D class averages exhibiting clear β-sheet separation. One round of 3D classification with a single class requested, a regularization parameter T of 3, and applied C2 helical symmetry was first used for broad alignment. The resulting volume did not show strand separation in the β-sheet and was low-pass filtered to 15 Å, then 10 Å, for two subsequent rounds of 3D auto-refinement, without optimization of helical rise and twist. We noticed some residual misalignment and performed 3D classification, unmasked, requesting 5 classes (K=5, T=2) using the newly implemented local averaging algorithm ^29^, implemented in RELION, selecting the best class with 73,287 particles. The particle stack was used for two successive auto-refinement procedures, the first unmasked and with local averaging, the second masked, low-passed filtered at 8 Å, and without local averaging. C2 symmetry was applied and allowed to modulate. The resolution at this stage reached around 4 Å with a B-factor estimated at −32 Å^2^ and β-strand separation. We then performed Bayesian polishing followed by multiple auto-refinement trials, allowing helical rise and twist optimization. Although resolutions between 3.5 and 4 Å were reached, symmetry parameters consistently converged poorly around 4.7 Å and 0.54°. We noticed some overfitting through residual signal in the phase-randomized FSC, particularly around 4.7 Å, and attributed this to incorrect symmetry parameters.

The resolution at this stage reached 3.5 Å, at which point we noticed similarities with a volume published prior that reached higher resolution with less over-fitting using a 2n-start symmetry operation. We therefore re-extracted the particles and performed auto-refinement procedures with a rise of 2.45 Å and twist of −179.66°. To note, reconstruction thus far had assumed right-handedness and we therefore modified the twist to reflect this. Two subsequent rounds of auto-refinements with decreasing low-pass filtering of the reference led to a resolution of 3.6 Å and no abnormal FSC. We followed with Bayesian polishing, auto-refinement, CTF corrections (third-order aberrations then per-particle defocus fitting), auto-refinement, and CTF corrections again (third-order aberrations then per-particle defocus fitting). Another round of Bayesian polishing was performed, this time omitting the last 20 frames as large motion was uncovered in the previous rounds, followed by auto-refinement. A final round of CTF corrections was performed in three steps: first estimating (anisotropic) magnification, then third- and -fourth order aberrations, then per-particle defocus and per-micrograph astigmatism. We found that estimating per-micrograph B-factor introduced errors and led to abnormal FSC curves. A final reconstruction was sharpened using standard post-processing procedure in RELION. An overall resolution of 3.1 Å was calculated from an FSC of 0.143 between the two independently refined maps.

#### Map sharpening and model building

Post-refinement map sharpening was performed using deepEMhancer ^65^ on a half-map from the final auto-refinement run. Here, we use manual measurement instead of automatic estimation for normalization parameters.

The inner layers of the resulting map were assessed to be similar to the ones found in Ghosh *et al*. ^29^. In addition, we were able to fit the two large residues pairs, I31/I32 and F19/F20 into large sidechain densities observed in the final volume. As such, these layers were initially built using 6W0O monomers in UCSF Chimera ^66^ and arranged in Coot ^67^. The densities in the inner layers of population B did not fit an entire monomer and we could only observe, and therefore place, densities for residues 15 to 39. The model was submitted to multiple rounds of real-space refinements in Phenix, with some re-arrangement in Coot ^67^ and ISOLDE ^69^ to alleviate clashes. MolProbity ^70^ and EMRinger ^71^ were used for model validation.

The outer layers of the map exhibit undefined densities. As described in the main text, these can potentially be attributed to β-hairpins layering upon the inner layers. Only segments K16–D23 and K28–M35 were modeled using the structure of *in vitro* Aβ40 exhibiting the canonical intra-molecular hydrophobic fold (PDB: 2LMN). Atoms beyond segments 16–23 and 29–35 were removed, the resulting disconnected molecule was arranged in Coot, then expanded and added to the inner layer initial model in UCSF Chimera in a parallel fashion. The initial model was subjected to multiple rounds of real-space refinement in Phenix, interspersed by arrangement in Coot and ISOLDE to resolve Ramachandran and sidechain outliers and alleviate clashes. The lack of definition in the outer densities led to much freedom of movement for the β-hairpins such that residues in these layers could not converge towards a stable conformation. While we are confident in the conformation of the inner layers, data contained in this work does not unequivocally support the presence of outer hairpins. These groups were therefore omitted from the deposited model.

## EXTENDED DATA

**Figure S1.**
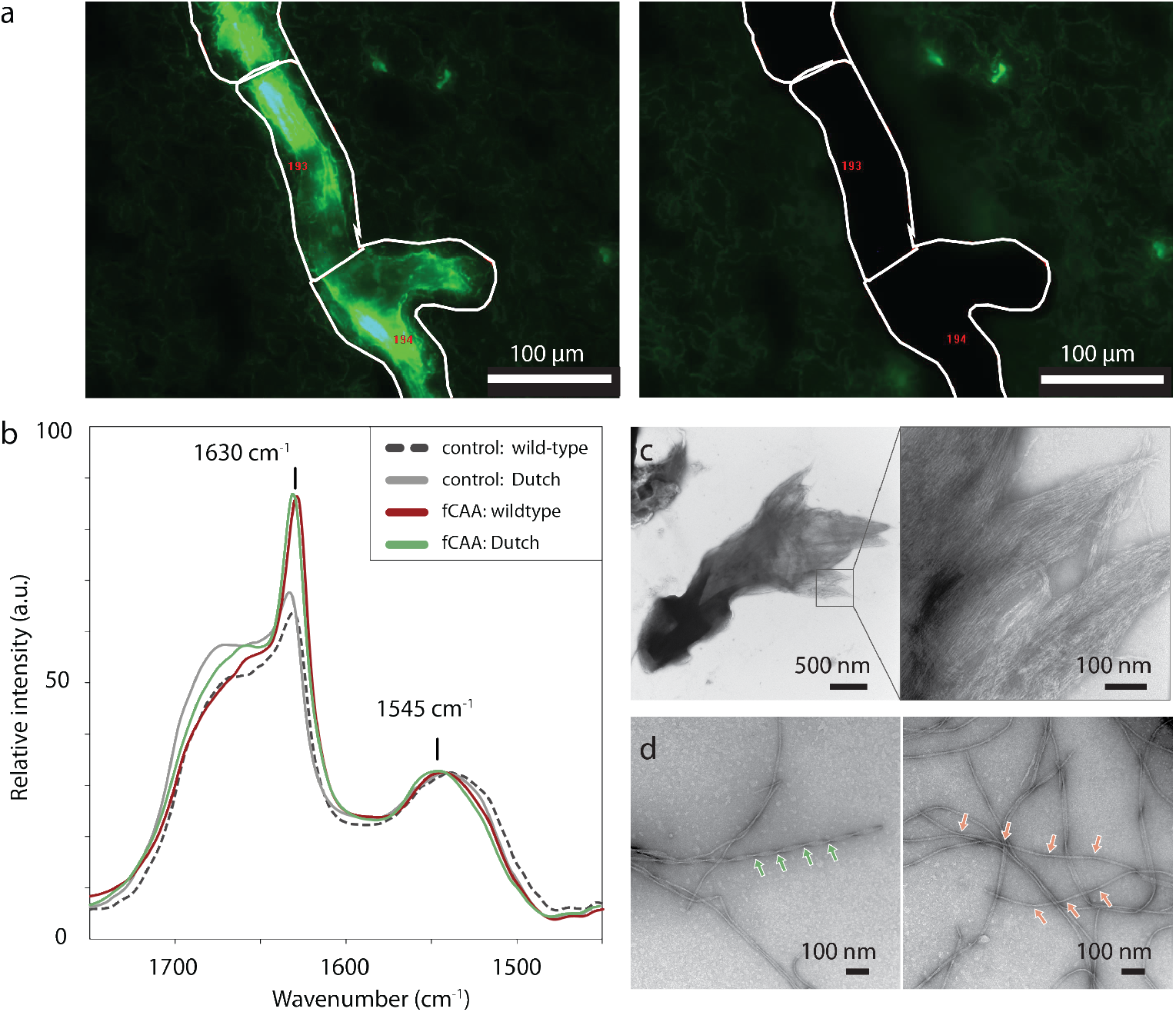
Templated growth of Aβ fibrils from fCAA-Dutch patient vascular deposits. (a) Representative fluorescence micrograph prior to (left) and following (right) laser capture microdissection of vascular amyloid from tissue slices of the fCAA-Dutch patient. The Aβ fibrils were labeled with thioflavin S (green). (b) FTIR spectroscopy confirmation of efficient templated growth from the brain-derived vascular deposits. FTIR spectroscopy provides a measure of fibril formation via a marked increase in the β-sheet–specific ~1630 cm^-1^ absorbance band and a shift of the amide band to ~1545-50 cm^-1^. The fibrils used for cryo-EM and NMR spectroscopy were generated using wild-type Aβ40 synthetic peptide (red trace) or synthetic Aβ40-Dutch peptide (green trace). Non-seeded negative controls of Aβ40 wild-type (dotted black trace) and Aβ40-Dutch (grey trace) monomers show a lack of increase at 1630 cm^-1^ and no shift at 1550 cm^-1^, indicative of non-fibrillar samples. (c) Representative negative stain TEM micrograph of disaggregated, proteinase K-treated, amyloid deposits isolated from the fCAA-Dutch patient using laser capture microdissection. Electron-dense aggregates (left) are composed of laterally associated fibrils (right). (d) Representative negative stain TEM micrographs of seeded fibrils at generation 3, after 3 rounds of templated growth. Both highly twisted (green arrow, population A) and slightly twisted (red arrows, population B) fibril morphologies are present, consistent with subsequent observations in cryo-EM micrographs.

**Figure S2.**
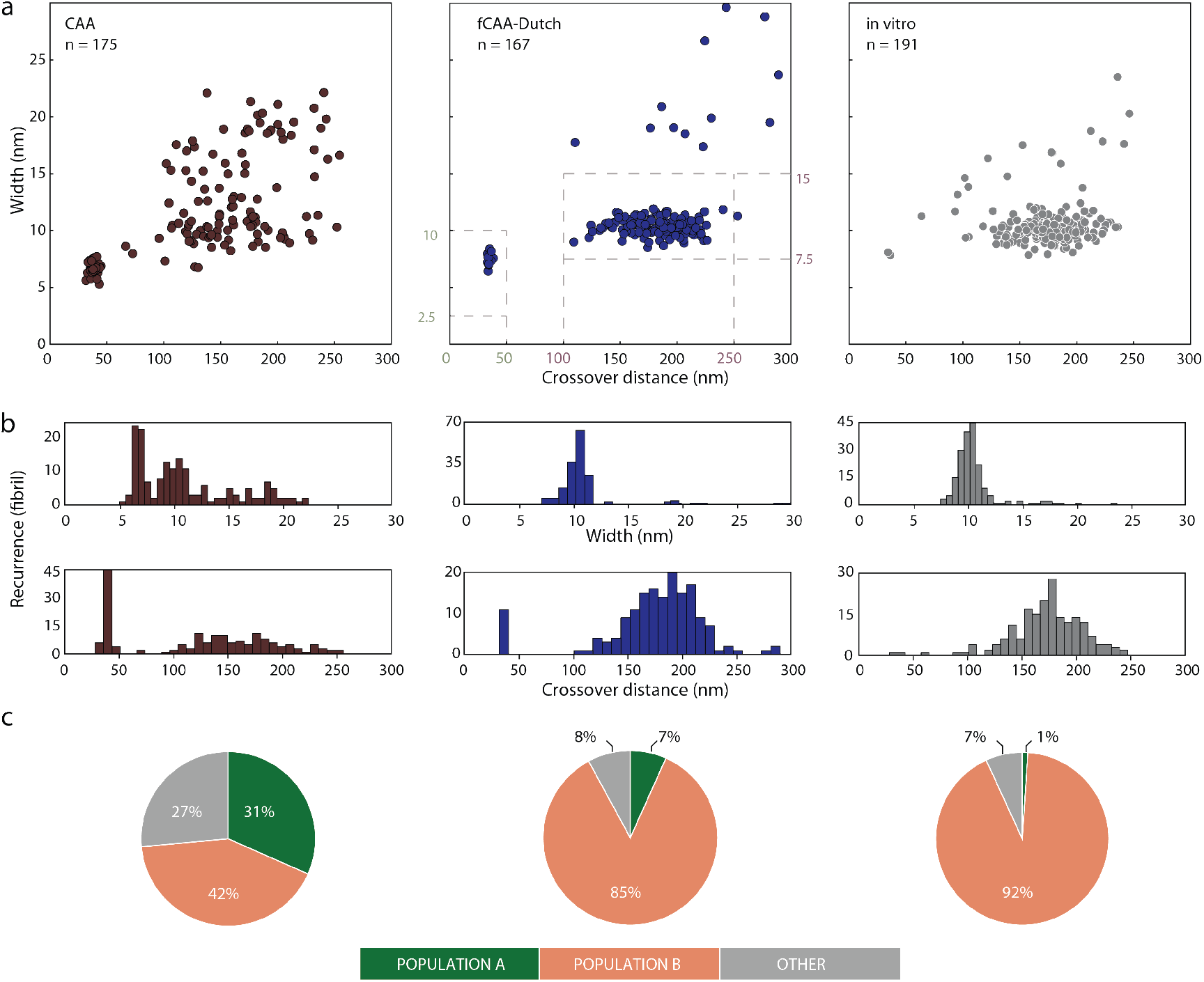
Two primary fibril populations can be distinguished by their helical morphology. (a) Expanded morphology summary plots for brain-derived CAA and fCAA-Dutch fibrils. An *in vitro* sample of Aβ40 wild-type fibrils was prepared for comparison by incubating Aβ40 under quiescent conditions. We find that most fibrils across all samples form two clusters with helical parameters highlighted by dashed lines in the fCAA-Dutch panel and classified as populations A and B, with corresponding ranges of crossover distance and fibril width highlighted. Population A adopts a highly twisted and narrow fibril architecture. Population B, however, exhibits high degrees of variability in all samples. (b) Crossover distance and fibril width are shown individually as histograms. The emergence of two populations is highlighted by the bimodal distribution of both crossover distances and width in some samples. Population A was not observed in the *in vitro* sample, suggesting it is unique to brain-derived fibrils. (c) Pie charts summarizing the fibril count in each population per patient sample indicate a net difference between the patients.

**Figure S3.**
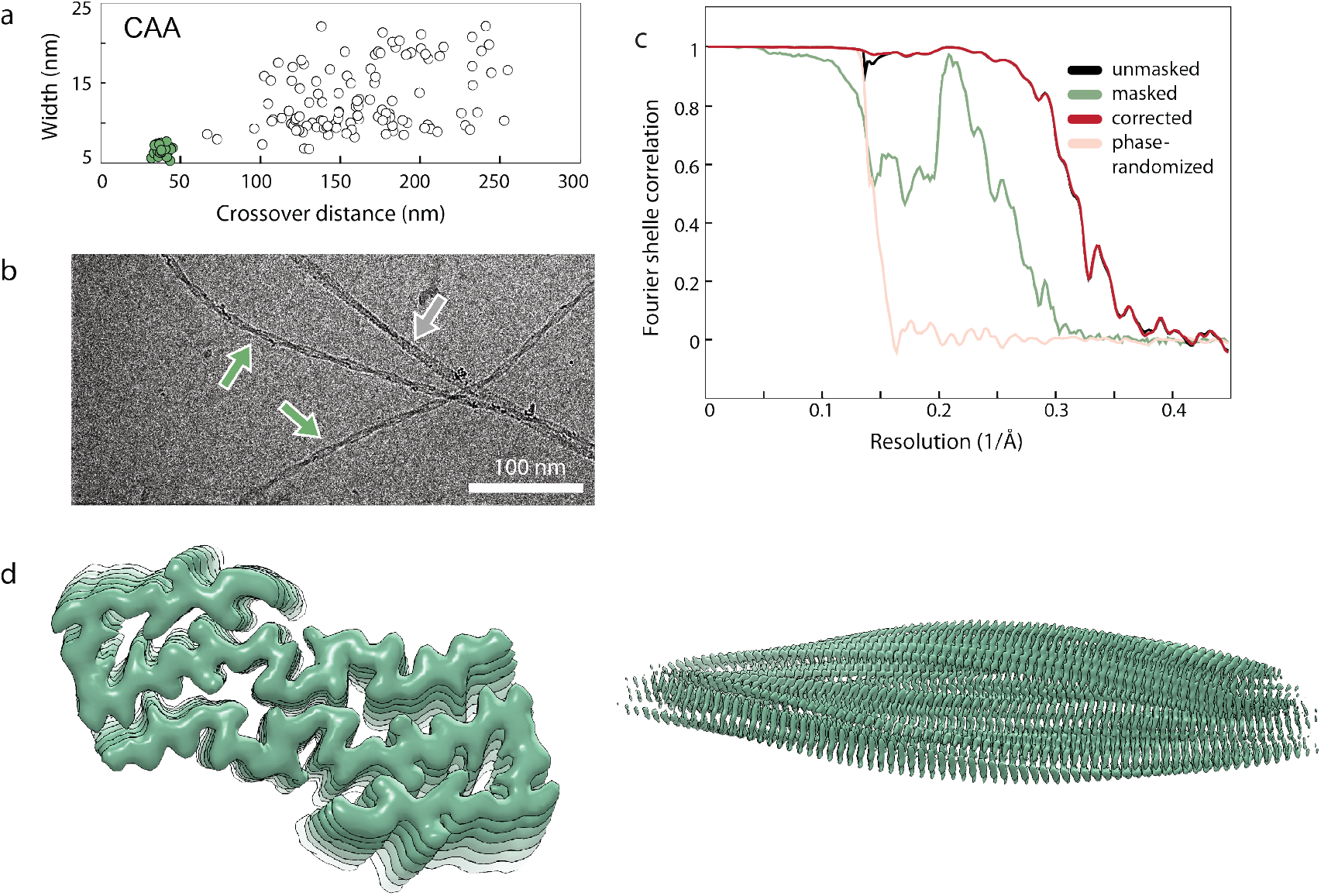
Details on population A in the CAA patient sample. (a) Morphology summary plot, shown in the main text, for this patient reveals a predominant population (green dots) with unique helical characteristics. The crossover distance averaging around 45 nm is not seen in *in vitro* fibrils, supporting the clinical relevance of this fibril population. (b) Representative cryo-EM micrograph highlighting fibril populations present in CAA sample, with the green arrows indicating the predominant population A targeted for helical reconstruction. (c) The quality of the curated particles and their correct alignment produced a resolution of 2.87 Å as estimated by a Fourier shell correlation with a threshold at 0.143. Phase-randomization procedures (red trace) do not indicate underlying issues with overfitting. Procedural details and further information are outlined in the Methods and in Table 1. (d) Top view and side view of the resulting sharpened map used for model-fitting. Visible β-strand separation and the appearance of β- and γ-carbons are visual indicators of a well-aligned high-resolution result.

**Figure S4.**
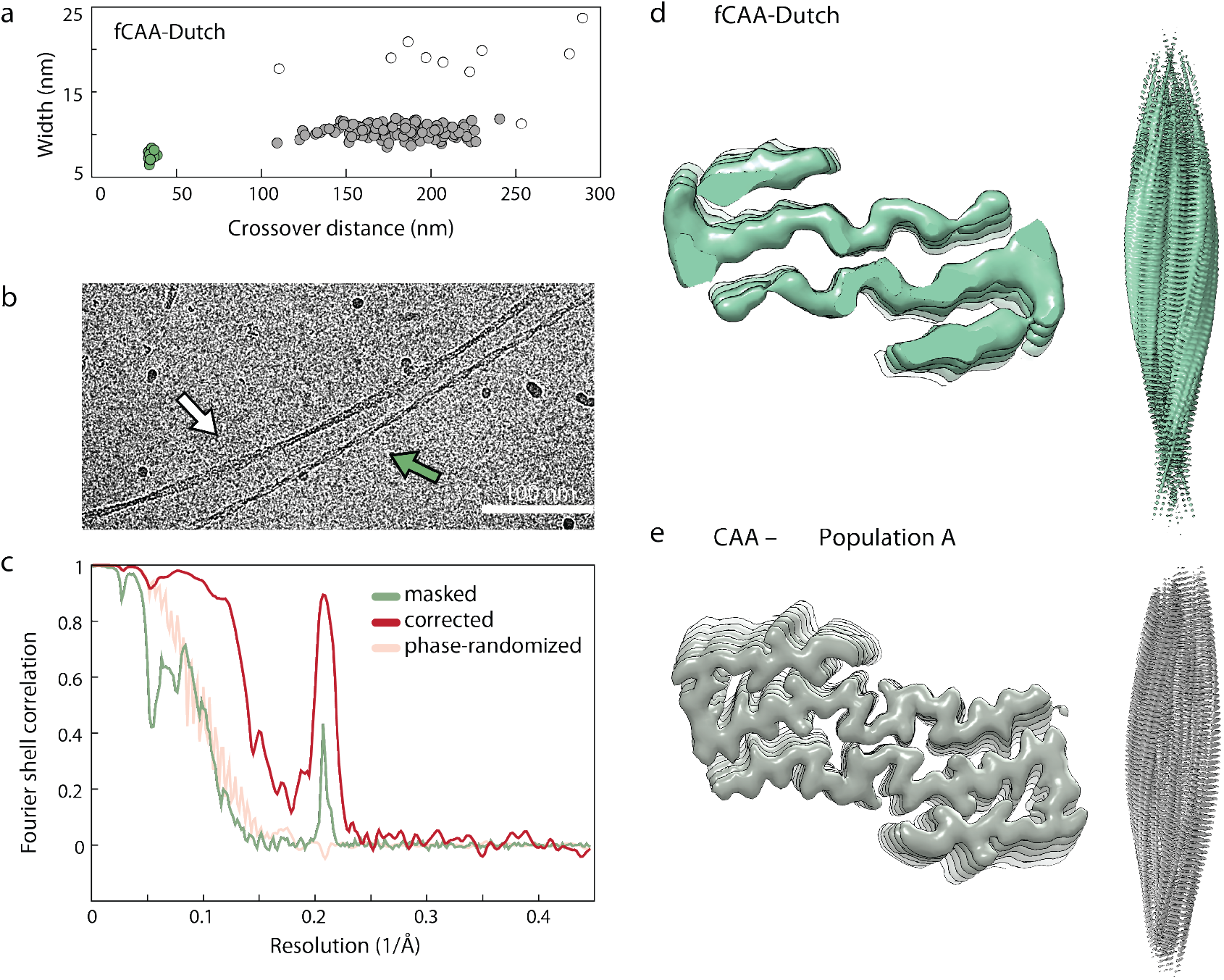
Population A is present in the vascular deposits from the fCAA-Dutch patient. (a) Morphology summary plot, shown in the main text, for the fCAA-Dutch patient shows a minor population (green dots) with identical helical characteristics as those present in the CAA patient from which we obtained a high-resolution structure. Grey dots represent the major fibril population (population B). (b) Representative cryo-EM micrograph highlighting both fibril populations present in the sample. We attempted helical reconstruction on the highly twisted fibrils (green arrows) as described in the Methods. White arrow points to a fibril from the major population B, corresponding to grey dots in panel a. (c) FSC curves for helical reconstruction of fCAA-Dutch fibrils. While the low number of particles precluded high-resolution reconstruction, we reached a resolution of about 7 Å, shown in the FSC curves. An increased Fourier-shell correlation between the half-maps around 4.7 Å reflects the expected bias introduced by a minimally low-pass filtered initial model. Residual signal past 10 Å in the phaserandomization procedure (light red trace) indicates some residual overfitting. Procedural details and further information are outlined in the Methods and in Table 1. (d and e) Comparison of the high-resolution cryo-EM density from the CAA patient (e) with the cryo-EM density obtained from the highly twisted fibrils in the fCAA-Dutch patient (d). The former was used as a low-passed initial model, successfully resulting in a similar backbone trace, albeit of lower resolution.

**Figure S5.**
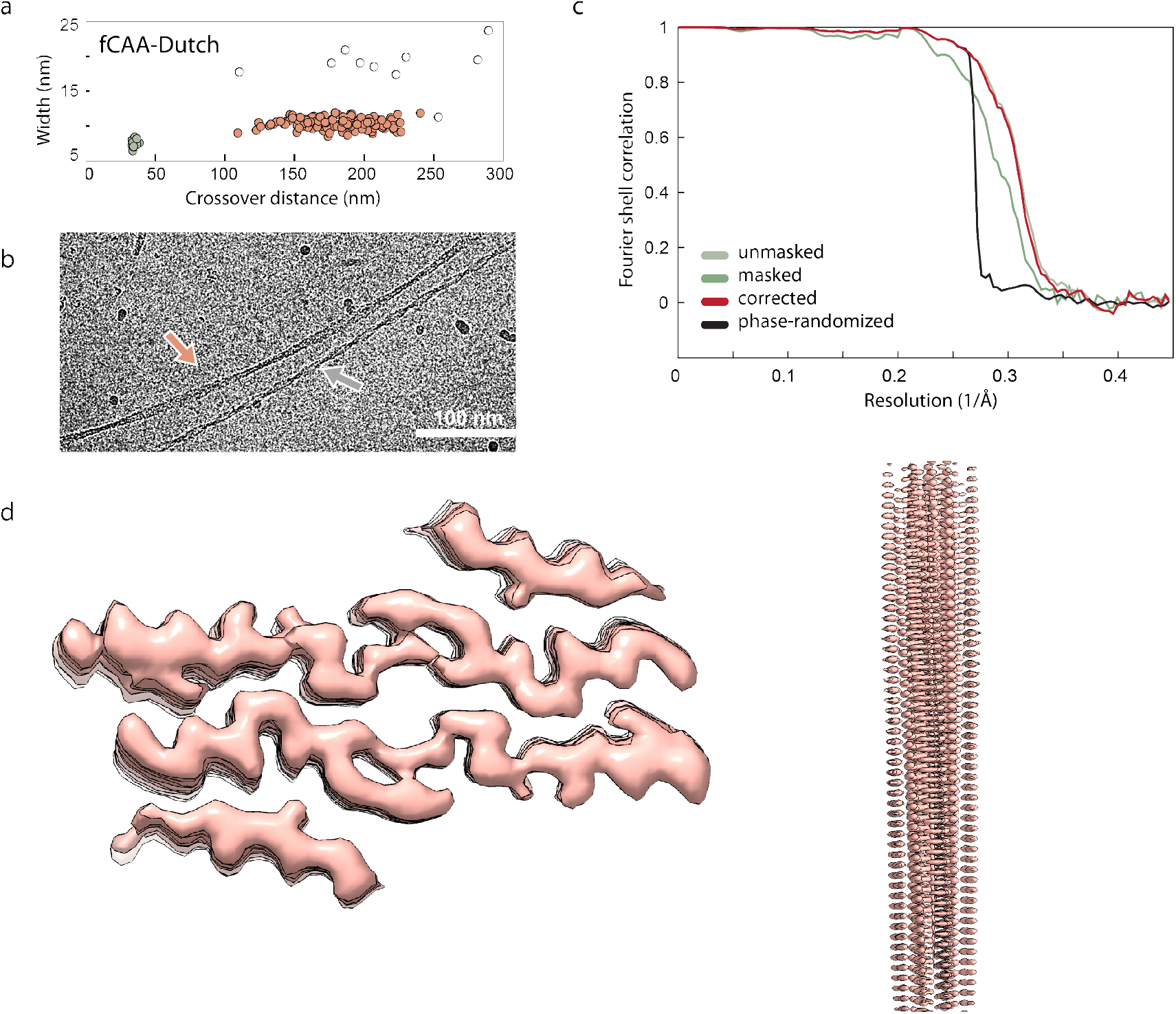
Details on population B in the fCAA-Dutch patient sample. (a) Morphology summary plot, shown in the main text, for the fCAA-Dutch patient reveals a predominant population (red dots) with helical characteristics reminiscent of commonly observed amyloid fibrils. Grey dots correspond to the minor population A. (b) Representative cryo-EM micrograph highlighting fibril populations present in the sample with red arrows indicating the predominant population B targeted for helical reconstruction. White arrow points to a fibril from the minor population A, corresponding to grey dots in panel a. (c) The final volume for this fibril population reached 3.09 Å as estimated by a Fourier shell correlation with a threshold of 0.143. Phase-randomization procedures (black trace) indicate very slight overfitting in the β-strand alignments. Procedural details and further information are outlined in the Methods and Table 1. (d) Top and side views of the resulting sharpened map used for model-fitting. Visible β-strand with no divergence in the half-maps (data not shown) indicates robust symmetry parameters. β-carbons and large sidechains are visible and allow for backbone tracing. The outer densities remained not well defined despite attempts at specifically targeting those areas. Ghosh et al. (2021) assigned this density to β-hairpin structures associated with the inner core of the fibril.

**Figure S6.**
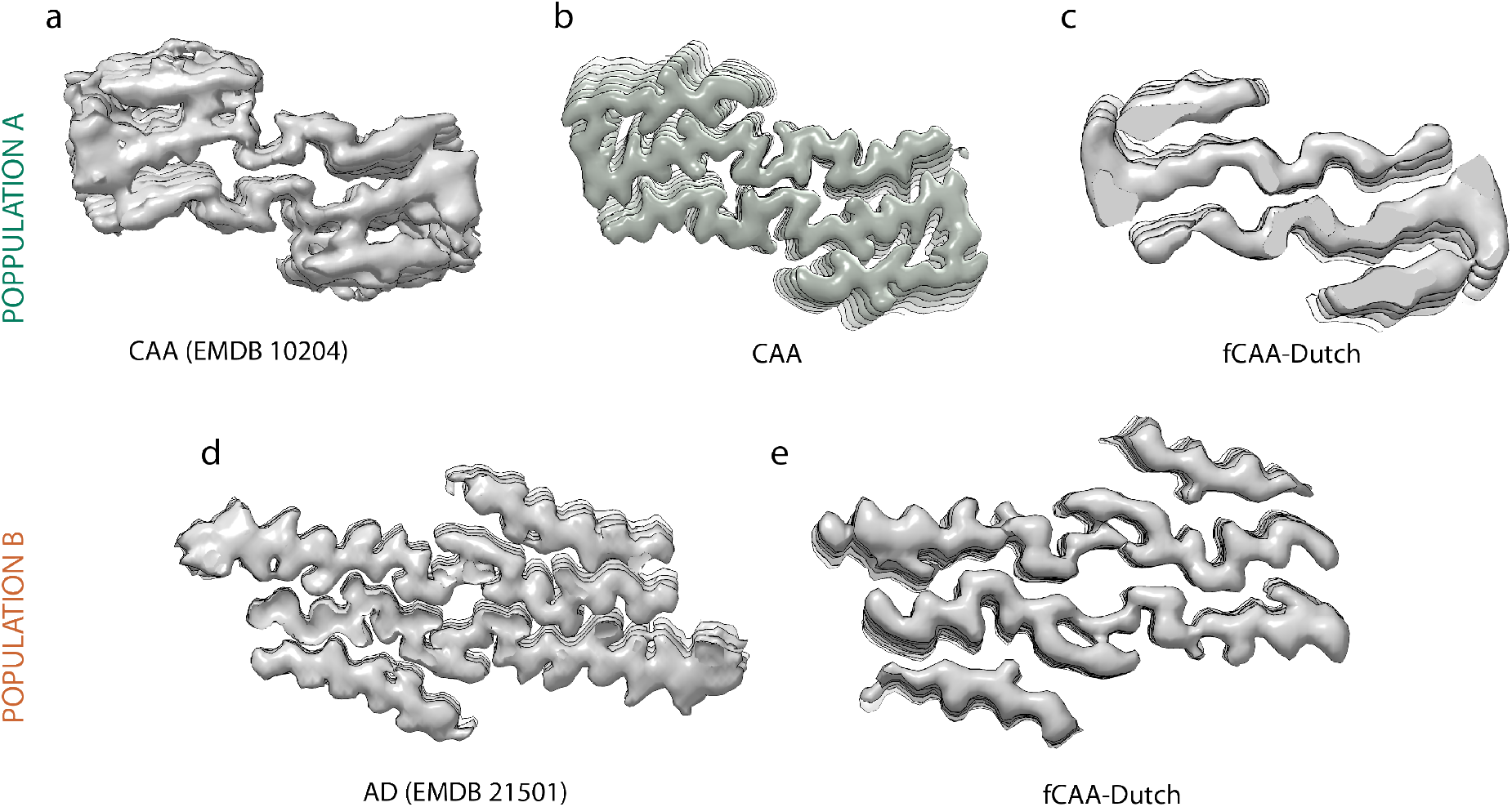
Comparison with published cryo-EM results. Strong similarities of cryo-EM volumes exist not only between patient samples investigated in this study, but with results from previously published studies. As detailed in the main text, we found two primary populations with distinct structure populate the sporadic and fCAA patients. These two populations are similar to fibrils extracted from a CAA patient (a), in the case of population A (b,c), and to fibrils extracted from parenchymal deposits of an AD patient (d) in the case of population B (e). The volumes presented here for comparison were developed by (a) Kollmer *et al.^1^* (EMDB: 10204) and (d) Ghosh *et al.^2^* (EMDB: 21501).

**Figure S7.**
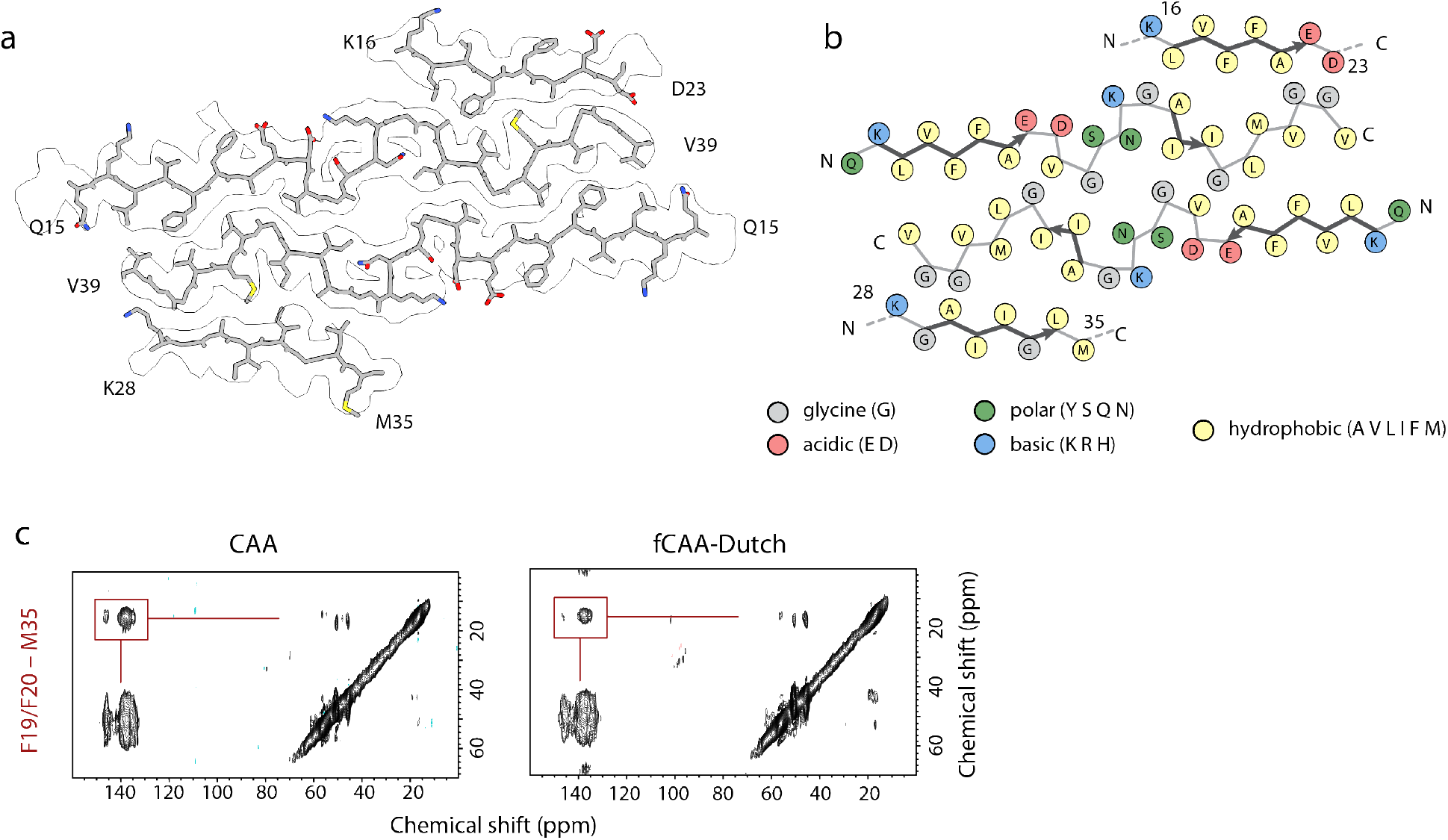
Potential packing interaction between F19 (outer layer) and M35 (inner layer) of population B. (a) Structure of population B in the outline of the cryo-EM density. The cryo-EM density is very similar to that obtained by Ghosh et al. (2021). In their study, they argued on the basis of solidstate NMR data that the outer densities resulted from associated β-hairpins. In our reconstructions, the outer densities of population B remained poorly defined. Nevertheless, we make note of a groove in the inner layers that would serve as a packing site for a large sidechain of the β-hairpin. On the basis of solid-state DARR NMR data, we suggest this large residue is F19 that packs against M35 in the inner layer. (b) Schematic representation of population B with each residue colored according to its electrostatic property, as indicated in the legend. The fold adopted by these β-hairpins is similar to the *in vitro* monomeric fold and mediated by the L17-A21 and A30-V36 hydrophobic segments, placing F19 near the middle of a β-strand. This results in the proximity of residues F19 and M35, an intermolecular contact otherwise absent from known structures, which we can use to interrogate the possibility of these β-hairpins layering atop fibrils in population B. (c) Two-dimensional ^13^C DARR NMR spectra of G4 fibrils templated off of brain-derived samples using the wild-type Aβ40 peptide containing 1–^13^C F4, 2–^13^C L17, ring–^13^C F19, ring–^13^C F20, 2–^13^C G33, 5–^13^C M35 and 1–^13^C G38. We observe a strong crosspeak for both patient samples (highlighted by a red box) between F19 and M35, consistent with the packing of F19 in the outer layer with M35 in the inner layer. The 2D ^13^C NMR spectra are shown with identical contour levels and were normalized manually.

**Figure S8.**
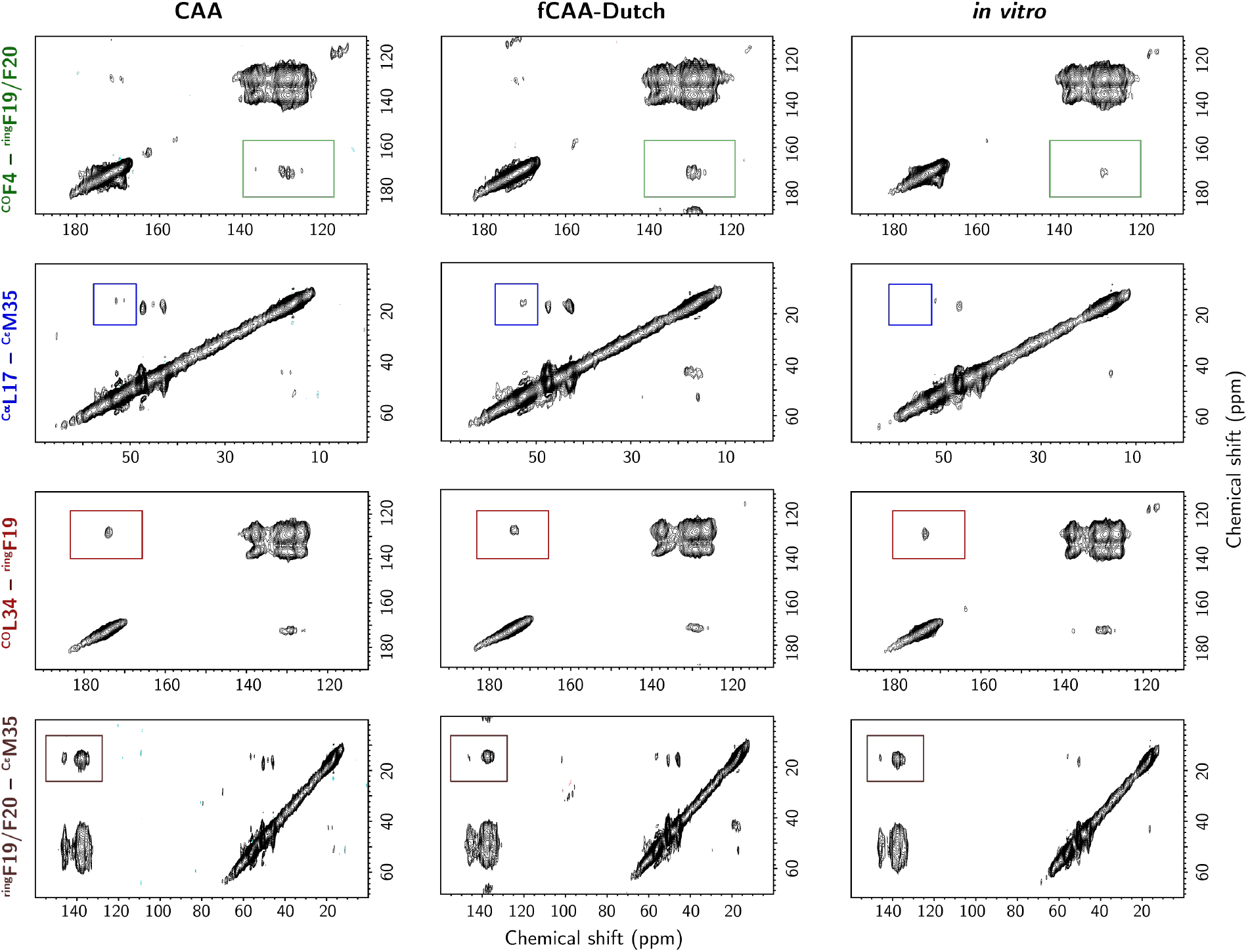
Solid-state NMR spectroscopy of CAA and fCAA-Dutch fibrils. Two-dimensional DARR ^13^C NMR spectra for residue pairs described in the main text are shown here for both patient samples. A portion of the generation 3 fibrils used for cryo-EM acquisition were used to produce ^13^C-labeled fibrils for NMR experiments via templated growth with wild-type Aβ40 peptide containing ring–^13^C F19 and U-^13^C L34 residues, or containing 1–^13^C F4, 2–^13^C L17, ring–^13^C F19, ring–^13^C F20, 2–^13^C G33, 5–^13^C M35 and 1–^13^C G38. Both labeling schemes were designed such that no cross-peak overlap and interpretation is unambiguous. Due to spinning side bands, the second labeling scheme required an NMR spectrum be acquired at two spinning speeds to counter cross-peak overlap.

## REFERENCES

1. Spina, S. et al. Comorbid neuropathological diagnoses in early versus late-onset Alzheimer’s disease. Brain 144, 2186–2198 (2021).

2. Toledo, J.B. et al. Contribution of cerebrovascular disease in autopsy confirmed neurodegenerative disease cases in the National Alzheimer’s Coordinating Centre. Brain 136, 2697–2706 (2013).

3. Sweeney, M.D. et al. Vascular dysfunction-The disregarded partner of Alzheimer’s disease. Alzheimers Dement 15, 158–167 (2019).

4. Reijmer, Y.D., van Veluw, S.J. & Greenberg, S.M. Ischemic brain injury in cerebral amyloid angiopathy. J. Cereb. Blood Flow Metab. 36, 40–54 (2016).

5. Greenberg, S.M. et al. Cerebral amyloid angiopathy and Alzheimer’s disease - one peptide, two pathways. Nature Rev. Neuro. 16, 30–42 (2020).

6. Wolfe, M.S. et al. Two transmembrane aspartates in presenilin-1 required for presenilin endoproteolysis and γ-secretase activity. Nature 398, 513–517 (1999).

7. Seino, Y. et al. Quantitative measurement of cerebrospinal fluid amyloid-β species by mass spectrometry. J. Alzheimers Dis. 79, 573–584 (2021).

8. Kummer, M.P. & Heneka, M.T. Truncated and modified amyloid-β species. Alzheimers Res. Ther. 6(2014).

9. Suzuki, N. et al. High tissue content of soluble Aβ1-40 is linked to cerebral amyloid angiopathy. Am. J. Pathol. 145, 452–460 (1994).

10. van Duinen, S.G. et al. Hereditary cerebral hemorrhage with amyloidosis in patients of Dutch origin is related to Alzheimer disease. Proc. Natl. Acad. Sci. USA 84, 5991–5994 (1987).

11. Van Broeckhoven, C. et al. Amyloid β protein precursor gene and hereditary cerebral hemorrhage with amyloidosis (Dutch). Science 248, 1120–1122 (1990).

12. Natté, R. et al. Dementia in hereditary cerebral hemorrhage with amyloidosis-Dutch type is associated with cerebral amyloid angiopathy but is independent of plaques and neurofibrillary tangles. Ann. Neurol. 50, 765–772 (2001).

13. Tycko, R. Physical and structural basis for polymorphism in amyloid fibrils. Protein Sci. 23, 1528–1539 (2014).

14. Cendrowska, U. et al. Unraveling the complexity of amyloid polymorphism using gold nanoparticles and cryo-EM. Proc. Natl. Acad. Sci. USA 117, 6866–6874 (2020).

15. Lu, J.-X. et al. Molecular structure of β-amyloid fibrils in Alzheimer’s disease brain tissue. Cell 154, 1257–68 (2013).

16. Qiang, W., Yau, W.M., Lu, J.X., Collinge, J. & Tycko, R. Structural variation in amyloid-β fibrils from Alzheimer’s disease clinical subtypes. Nature 541, 217–221 (2017).

17. Condello, C. et al. Structural heterogeneity and intersubject variability of Aβ in familial and sporadic Alzheimer’s disease. Proc. Natl. Acad. Sci. USA 115, E782–E791 (2018).

18. Rasmussen, J. et al. Amyloid polymorphisms constitute distinct clouds of conformational variants in different etiological subtypes of Alzheimer’s disease. Proc. Natl. Acad. Sci. USA 114, 13018–13023 (2017).

19. Liu, J.L. et al. Amyloid structure exhibits polymorphism on multiple length scales in human brain tissue. Sci. Rep. 6(2016).

20. Rutgers, K.S. et al. Differential recognition of vascular and parenchymal β amyloid deposition. Neuro. Aging 32, 1774–1783 (2011).

21. Han, B.H. et al. Resorufin analogs preferentially bind cerebrovascular amyloid: potential use as imaging ligands for cerebral amyloid angiopathy. Mol. Neurodegen. 6 (2011).

22. Schrag, M. et al. Effect of cerebral amyloid angiopathy on brain iron, copper, and zinc in Alzheimer’s disease. J. Alzheimers Dis. 24, 137–149 (2011).

23. Schweighauser, M. et al. Structures of alpha-synuclein filaments from multiple system atrophy. Nature 585, 464–469 (2020).

24. Kollmer, M. et al. Cryo-EM structure and polymorphism of Aβ amyloid fibrils purified from Alzheimer’s brain tissue. Nature Comm. 10, DOI: 10.1038/s41467-019-12683-8 (2019).

25. Wickramasinghe, A. et al. Sensitivity-enhanced solid-state NMR detection of structural differences and unique polymorphs in pico- to nanomolar amounts of brain-derived and synthetic 42-residue amyloid-β fibrils. J. Am. Chem. Soc. 143, 11462–11472 (2021).

26. Ghosh, U., Yau, W.M., Collinge, J. & Tycko, R. Structural differences in amyloid-β fibrils from brains of nondemented elderly individuals and Alzheimer’s disease patients. Proc. Natl. Acad. Sci. USA 118(2021).

27. Yang, Y. et al. Cryo-EM structures of amyloid-β 42 filaments from human brains. Science 375, 167–172 (2022).

28. Greenberg, S.M. et al. Outcome markers for clinical trials in cerebral amyloid angiopathy. Lancet Neurology 13, 419–428 (2014).

29. Ghosh, U., Thurber, K.R., Yau, W.M. & Tycko, R. Molecular structure of a prevalent amyloid-β fibril polymorph from Alzheimer’s disease brain tissue. Proc. Natl. Acad. Sci. USA 118, DOI: 10.1073/pnas.2023089118 (2021).

30. Irizarry, B.A. et al. Human cerebral vascular amyloid contains both anti-parallel and parallel in-register Aβ40 fibrils. J. Biol. Chem. 297, 101259–101272 (2021).

31. Ghosh, U., Yau, W.M. & Tycko, R. Coexisting order and disorder within a common 40-residue amyloid-β fibril structure in Alzheimer’s disease brain tissue. Chem. Commun. 54, 5070–5073 (2018).

32. Hu, Y. et al. β-Sheet structure within the extracellular domain of C99 regulates amyloidogenic processing. Sci. Rep. 7, 17159–17166 (2017).

33. Kukar, T.L. et al. Lysine 624 of the amyloid precursor protein (APP) is a critical determinant of amyloid β peptide length: Support for a sequential model of γ-secretase intramembrane proteolysis and regulation by the amyloid β precursor protein (APP) juxtamembrane region. J. Biol. Chem. 286, 39804–39812 (2011).

34. Petit, D. et al. Extracellular interface between APP and nicastrin regulates Aβ length and response to gamma-secretase modulators. EMBO J 38(2019).

35. Mikros, E. et al. High-resolution NMR spectroscopy of the β-amyloid(1-28) fibril typical for Alzheimer’s disease. Angew. Chem.-Int. Edit. 40, 3603–3605 (2001).

36. Fraser, P.E., Nguyen, J.T., Surewicz, W.K. & Kirschner, D.A. pH-Dependent structural transitions of Alzheimer amyloid peptides. Biophys. J. 60, 1190–1201 (1991).

37. Alvarez, X.A., MiguelHidalgo, J.J., FernandezNovoa, L. & Cacabelos, R. Intrahippocampal injections of the β-amyloid 1-28 fragment induces behavioral deficits in rats. Methods Find. Exp. Clin. Pharmacol. 19, 471–479 (1997).

38. Jonsson, T. et al. A mutation in APP protects against Alzheimer’s disease and age-related cognitive decline. Nature 488, 96–99 (2012).

39. Di Fede, G. et al. A recessive mutation in the APP gene with dominant-negative effect on amyloidogenesis. Science 323, 1473–1477 (2009).

40. Nguyen, P.H., Tarus, B. & Derreumaux, P. Familial Alzheimer A2V mutation reduces the intrinsic disorder and completely changes the free energy landscape of the Aβ1-28 monomer. J. Phys. Chem. B 118, 501–510 (2014).

41. Brinkmalm, G. et al. Identification of neurotoxic cross-linked amyloid-β dimers in the Alzheimer’s brain. Brain 142, 1441–1457 (2019).

42. Wilhelmus, M.M.M., Bol, J., van Duinen, S.G. & Drukarch, B. Extracellular matrix modulator lysyl oxidase colocalizes with amyloid-β pathology in Alzheimer’s disease and hereditary cerebral hemorrhage with amyloidosis-Dutch type. Exp. Gerontol. 48, 109–114 (2013).

43. Gilad, G.M., Kagan, H.M. & Gilad, V.H. Lysyl oxidase, the extracellular matrix-forming enzyme, in rat brain injury sites. Neurosci. Lett. 310, 45–48 (2001).

44. Paravastu, A.K., Leapman, R.D., Yau, W.M. & Tycko, R. Molecular structural basis for polymorphism in Alzheimer’s β-amyloid fibrils. Proc Natl Acad Sci U S A 105, 18349–54 (2008).

45. Qiang, W., Yau, W.-M., Luo, Y., Mattson, M.P. & Tycko, R. Antiparallel β-sheet architecture in Iowa-mutant β-amyloid fibrils. Proc. Natl. Acad. Sci. USA 109, 4443–4448 (2012).

46. Maji, S.K. et al. Amino acid position-specific contributions to amyloid β-protein oligomerization. J. Biol. Chem. 284, 23580–23591 (2009).

47. Kumar-Singh, S. Hereditary and sporadic forms of Aβ-cerebrovascular amyloidosis and relevant transgenic mouse models. Int. J. Mol. Sci. 10, 1872–1895 (2009).

48. Bugiani, O. et al. Hereditary cerebral hemorrhage with amyloidosis associated with the E693K mutation of APP. Arch. Neurol. 67, 987–995 (2010).

49. Sandberg, A. et al. Stabilization of neurotoxic Alzheimer amyloid-β oligomers by protein engineering. Proc. Natl. Acad. Sci. USA 107, 15595–15600 (2010).

50. Rajpoot, J. et al. Insights into cerebral amyloid angiopathy type 1 and type 2 from comparisons of the fibrillar assembly and stability of the Aβ40-Iowa and Aβ40-Dutch peptides. Biochemistry 61, 1181–1198 (2022).

51. Tornquist, M. et al. Secondary nucleation in amyloid formation. Chem. Commun. 54, 8667–8684 (2018).

52. Crooks, E.J. et al. Copper stabilizes antiparallel β-sheet fibrils of the amyloid-β40 (Aβ40)-Iowa variant. J. Biol. Chem. 295, 8914–8927 (2020).

53. Hondius, D.C. et al. Proteomics analysis identifies new markers associated with capillary cerebral amyloid angiopathy in Alzheimer’s disease. Acta Neuro. Comm. 6(2018).

54. Cao, Q., Boyer, D.R., Sawaya, M.R., Ge, P. & Eisenberg, D.S. Cryo-EM structures of four polymorphic TDP-43 amyloid cores. Nat. Struct. Mol. Biol. 26, 619–627 (2019).

55. Heise, H. et al. Molecular-level secondary structure, polymorphism, and dynamics of full-length α-synuclein fibrils studied by solid-state NMR. Proc. Natl. Acad. Sci. USA 102, 15871–15876 (2005).

56. Scherpelz, K.P. et al. Atomic-level differences between brain parenchymal- and cerebrovascular-seeded Aβ fibrils. Sci. Rep. 11 (2021).

57. Bakrania, P. et al. Discovery of a novel pseudo β-hairpin structure of N-truncated amyloid-β for use as a vaccine against Alzheimer’s disease. Mol. Psychiatry.

58. Arndt, J.W. et al. Structural and kinetic basis for the selectivity of aducanumab for aggregated forms of amyloid-β. Sci. Rep. 8(2018).

59. Bohrmann, B. et al. Gantenerumab: a novel human anti-Aβ antibody demonstrates sustained cerebral amyloid-β binding and elicits cell-mediated removal of human amyloid-β. J. Alzheimers Dis. 28, 49–69 (2012).

60. van Dyck, C.H. et al. Lecanemab in early Alzheimer’s disease. New Engl. J. Med. (2022).

## References - Cited in On-Line Methods Only

61. Kremer, J.R., Mastronarde, D.N. & McIntosh, J.R. Computer visualization of three-dimensional image data using IMOD. J. Struct. Biol. 116, 71–76 (1996).

62. Zheng, S.Q. et al. MotionCor2: anisotropic correction of beam-induced motion for improved cryo-electron microscopy. Nat. Methods 14, 331–332 (2017).

63. Zhang, K. Gctf: Real-time CTF determination and correction. J. Struct. Biol. 193, 1–12 (2016).

64. Scheres, S.H.W. Amyloid structure determination in RELION-3.1. Acta Crys. Section D Struct. Biol. 76, 94–101 (2020).

65. Sanchez-Garcia, R. et al. DeepEMhancer: a deep learning solution for cryo-EM volume postprocessing. Commun. Biol. 4 (2021).

66. Goddard, T.D., Huang, C.C. & Ferrin, T.E. Visualizing density maps with UCSF Chimera. J. Struct. Biol. 157, 281–287 (2007).

67. Emsley, P., Lohkamp, B., Scott, W.G. & Cowtan, K. Features and development of Coot. Acta Crys. Section D Struct. Biol. 66, 486–501 (2010).

68. Liebschner, D. et al. Macromolecular structure determination using X-rays, neutrons and electrons: recent developments in Phenix. Acta Crys. Section D Struct. Biol. 75, 861–877 (2019).

69. Croll, T.I. ISOLDE: a physically realistic environment for model building into low-resolution electron-density maps. Acta Crys. Section D Struct. Biol. 74, 519–530 (2018).

70. Chen, V.B. et al. MolProbity: all-atom structure validation for macromolecular crystallography. Acta Crys. Section D Struct. Biol. 66, 12–21 (2010).

71. Barad, B.A. et al. EMRinger: side chain directed model and map validation for 3D cryo-electron microscopy. Nat. Methods 12, 943–946 (2015).

## References

1. Kollmer, M. et al. Cryo-EM structure and polymorphism of Abeta amyloid fibrils purified from Alzheimer’s brain tissue. Nat. Commun. 10, 4760 (2019).

2. Ghosh, U., Thurber, K.R., Yau, W.M. & Tycko, R. Molecular structure of a prevalent amyloidbeta fibril polymorph from Alzheimer’s disease brain tissue. Proc. Natl. Acad. Sci. USA 118, e2023089118 (2021).

3. Liebschner, D. et al. Macromolecular structure determination using X-rays, neutrons and electrons: recent developments in Phenix. Acta Crys. Section D Struct. Biol. 75, 861–877 (2019).

4. Barad, B.A. et al. EMRinger: side chain-directed model and map validation for 3D cryo-electron microscopy. Nat Methods 12, 943–6 (2015).

5. Emsley, P., Lohkamp, B., Scott, W.G. & Cowtan, K. Features and development of Coot. Acta Crys. Section D Struct. Biol. 66, 486–501 (2010).

